# In silico Antibody-Peptide Epitope prediction for Personalized cancer therapy

**DOI:** 10.1101/2023.01.23.525181

**Authors:** Ivan Jacobs, Lim Chwee Ming, Jamie Mong, Maragkoudakis Emmanoil, Nishant Malik

## Abstract

The human leukocyte antigen (HLA) system is a complex of genes on chromosome 6 in humans that encodes cell-surface proteins responsible for regulating the immune system. Viral peptides presented to cancer cell surfaces by the HLA trigger the immune system to kill the cells, creating Antibody-peptide epitopes (APE). This study proposes an in-silico approach to identify patient-specific APEs by applying complex networks diagnostics on a novel multiplex data structure as input for a deep learning model. The proposed analytical model identifies patient and tumor-specific APEs with as few as 20 labeled data points. Additionally, the proposed data structure employs complex network theory and other statistical approaches that can better explain and reduce the black box effect of deep learning. The proposed approach achieves an F1-score of 80% and 93% on patients one and two respectively and above 90% on tumor-specific tasks. Additionally, it minimizes the required training time and the number of parameters.

## 1 INTRODUCTION

The human leukocyte antigen (HLA) system or complex is a complex of genes on chromosome 6 in humans that encode cell-surface proteins responsible for regulating the immune system. The HLA system also known as the human version of the major histocompatibility complex (MHC) is found in many animals.

HLA genes are highly polymorphic, which means that there are thousands of different forms of these genes called alleles, allowing them to fine-tune the adaptive immune system. The proteins encoded by certain genes are also known as antigens, because of their historic discovery as factors in organ transplants.

As shown in Figure 1 HLA’s proteins present viral peptides from inside the cell to the surface of the cell. For example, if the cell is infected by a virus or is cancerous, the HLA system brings abnormal fragments, called peptides, to the surface of the cell so that the cell can be destroyed by the immune system.

**Figure 1.**
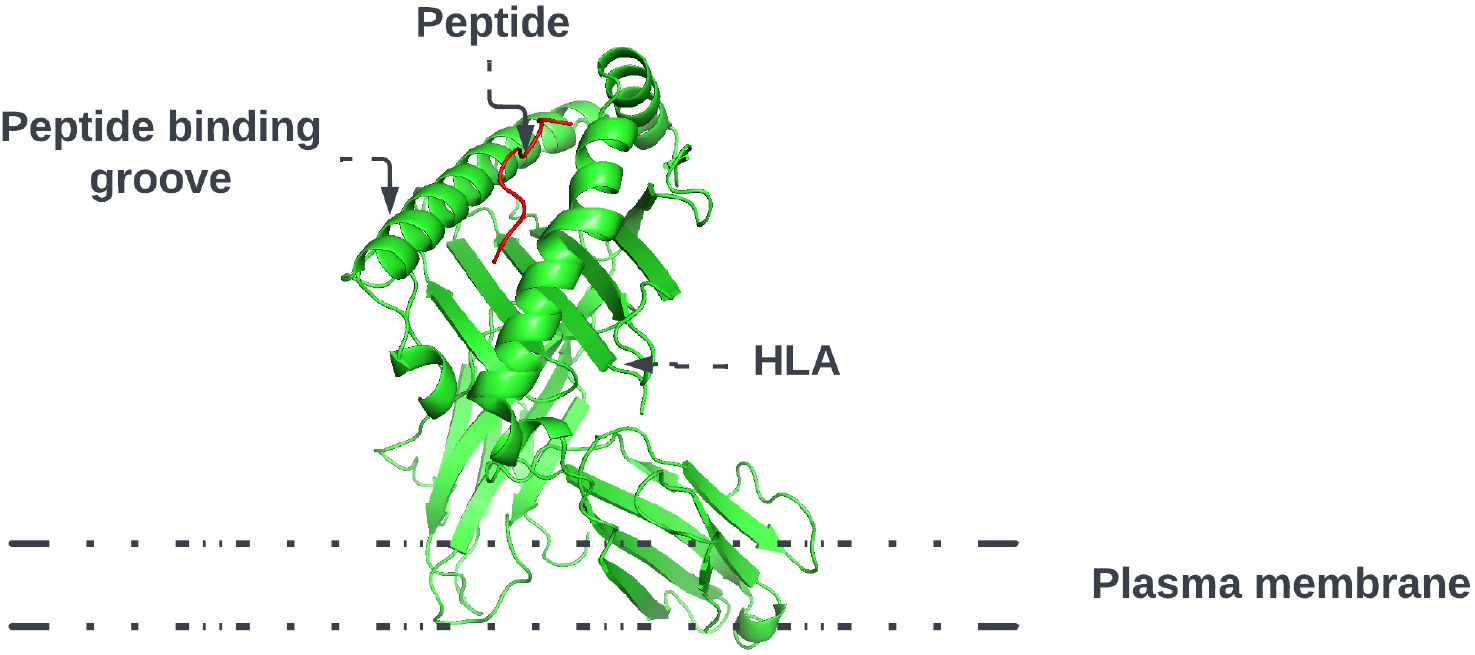
HLA proteins (green) display peptides (red) from inside the cell to help immune cells find cancerous or infected cells

Predicting the specific HLA peptide combination that will present the peptide to the cell’s surface permits the creation of a treatment that will trigger the human immune system to destroy the cell. Specifically, in cancer, this ability is essential, given that cancer is highly mutagenic with tumor and patient-specific mutations. This means that patients with the same tumor type will have different mutations that result in different reactions to the same treatment.

Advances in Deoxyribonucleic acid (DNA) sequencing, Messenger Ribonucleic acid (mRNA) vaccines, and high computational power allow us to work toward patient-specific therapy. This approach, called personalized mRNA-based antitumor vaccine, visualized in Figure 2, is bound to play a major role in the future.

**Figure 2.**
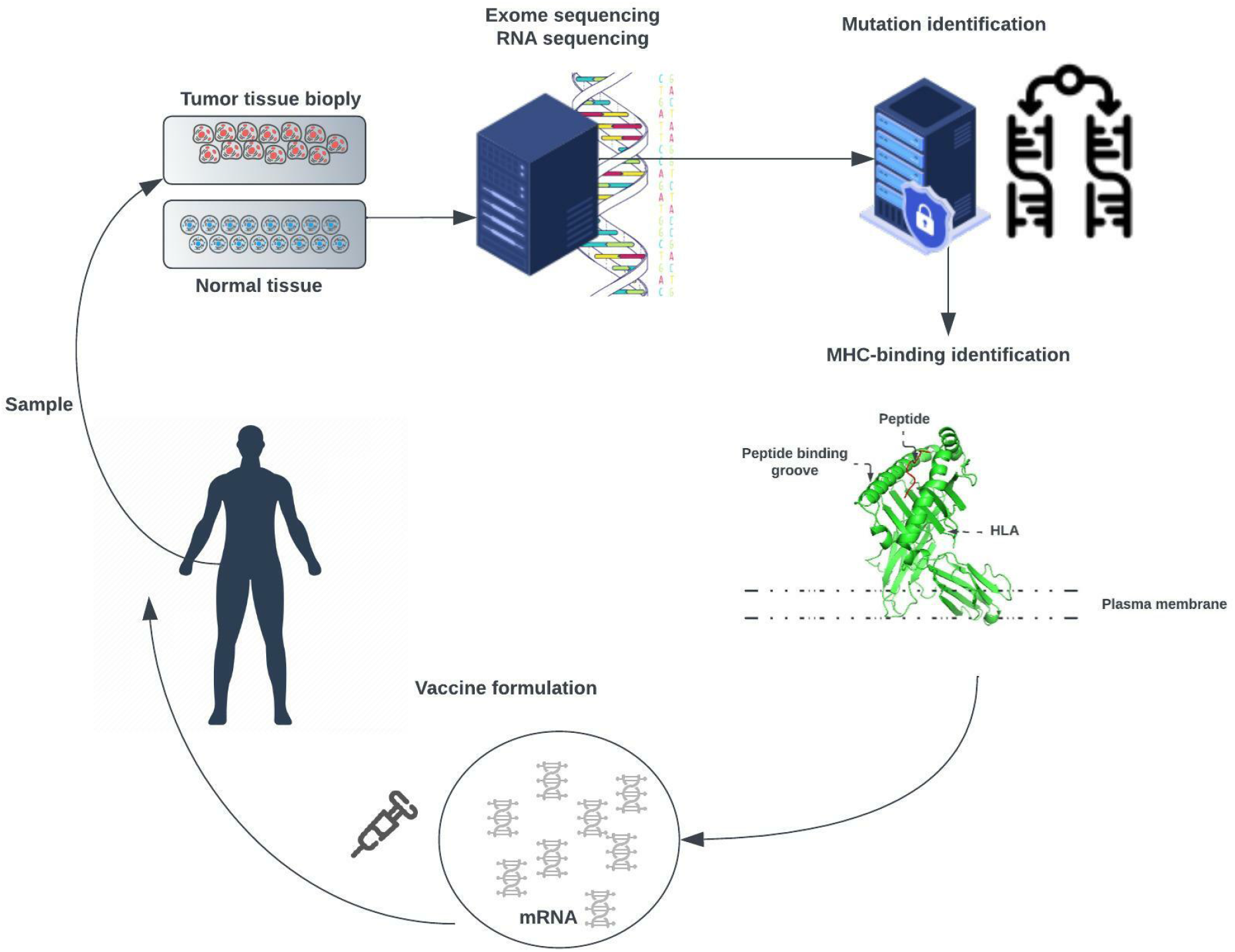
The exome of tumor cells isolated from a biopsy sample and the exome of normal cells are compared to identify tumor-specific mutations. Point non-synonymous mutations, gene deletions, or rearrangements can give rise to neoantigens. Several bioinformatics tools are used to predict major histocompatibility complex (MHC) class I and class II binding (necessary for recognition by T cells) and RNA expression presence of the mutated antigen among tumor cells (clonality). RNA sequencing enables verification that the gene encoding the neoantigen is actually expressed by tumor cells. A tandem gene encoding several neoantigen peptides is cloned into a plasmid and transcribed to mRNA. Finally, these mRNAs are injected as naked RNA, formulated into liposomes, or loaded into dendritic cells.

The approach is meant to trigger an antitumor immune response in patients by challenging them with mRNAs encoding tumor-specific antigens (1). These mRNAs can be directly injected as naked RNA or loaded into patient-derived dendritic cells.

In this work, we propose to extend the approach with additional laboratory and analytical optimization steps. Concretely DNA sequenced from the patient is used to select candidate peptides that will result from gene expression. As the space of possible combinations is huge a subset of potential peptides is synthesized and their reaction to the patient’s specific HLA alleles is tested by applying an enzyme-linked immunospot (ELISpot) assay. An enzyme-linked immunospot (ELISpot) assay (2), shown in Figure 3, is a highly versatile and sensitive technique that is used for qualitative and quantitative measurement of the cytokine-secreting cells at the single-cell level. (3)

**Figure 3.**
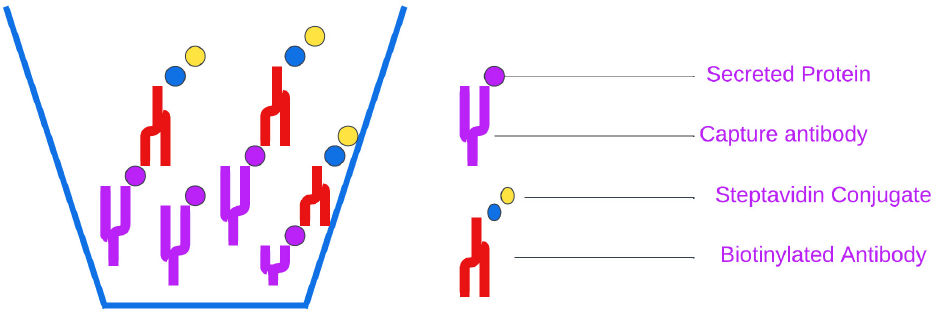
Visualisation of an enzyme-linked immunospot (ELISpot) Assay

This assay involves culturing cells on a surface on which a reagent (e.g., a specific capture antibody) is immobilized. Secreted proteins by the cells, such as cytokines, will be captured by the specific antibodies on the surface. Post-appropriate incubation time, cells are washed, and the secreted molecule is identified by applying a detection antibody. Adding a substrate produces a colored fluorescent or luminescent reaction (e.g., visible spots on the surface). Each spot corresponds to an individual cytokine-secreting cell and indicates the reactants’ concentration. The generated data permits fine-tuning an analytical approach to a specific patient’s and tumor’s mutations and reevaluating the peptide sequences with the purpose of selecting the optimal HLA allele-peptide combination. Once a subset of peptides is identified an mRNA vaccine is created that will force the body to trigger an immune response and destroy cancerous cells. The mRNA vaccine creation and evaluation are outside the scope of this work. The purpose of this work is to identify an analytical approach that can predict HLA-peptide interaction. With this effort, we hope to provide a framework that gives the ability to select and optimize personalized analytical methods from a wide range of possibilities as graph theory, machine learning, deep learning, meta-learning, and machine learning on graphs and leverage on high-performance computing with graphic processing unit (GPU) acceleration.

## 2 RELATED WORK

Epitope prediction is an important approach in tumor immunology and immunotherapy. The main classes of importance in the HLA molecules are the Class I and II molecules, which present epitopes to CD8^+^ T cells and CD4^+^ T cells respectively. Methods for assessing immunogenicity are MS-based MHC-I peptide binding prediction and immunogenicity verification by specific response assays (4, 5, 6). Many analytical approaches have been applied to distinguish two main types of approaches, allele-specific and pan-specific. Where the former trains one model for every MHC-I allele (7, 8, 9, 10, 11, 12) and the latter considers them to be as one and trains a global model together on both (13, 14, 15, 16). Methods with the highest achieved accuracy use data from The Immune Epitope Database (17). In recent years, the high rate of deep learning research resulted in a variety of deep learning based methods proposed by researchers (12, 13, 18, 19, 20). A number of approaches combine graph theory principles with deep learning or deep learning on graphs (21, 22, 23, 24) in order to detect interactive propensities embedded in HLA-peptide pairs.

Even though high affinity in an MHC-peptide complex tends to be associated with immune responsiveness, it is not sufficient to define immunogenicity. The existing analytical models lack several aspects influential to immunogenicities, such as the rate of expression, failure to represent sophisticated dynamics in molecular systems, and abundance of proteins. A number of top-ranking analytical approaches regularly falsely predict neoantigens (25, 26, 27). Hence, the need to better comprehend immune responsiveness based on MHC-peptide complex and dynamic structures and interaction in the context of a complex dynamical system environment is key for peptide-based personalized vaccines.

## 3 APPROACH

To compare and predict systems interactions and behavior we will look at measures and metrics that these networks express. Calculating and assigning these metrics for every individual system permits us to create a data set that can be used in statistical, machine, and deep learning analysis approaches. Our approach takes the network measures as input and makes a binary decision about whether a system composed of multiple patient-specific peptides and HLAs will result in the presentation of viral epitopes (in known EBV-driven cancer such as nasopharyngeal cancer) on the surface of the cancer cells. As shown in Figure 4, the work can be divided into the following phases: (1) Data Collection, (2) ELISpot assay, (3) Structures and Measurements generation, and (4) Analytical approach evaluation.

**Figure 4.**
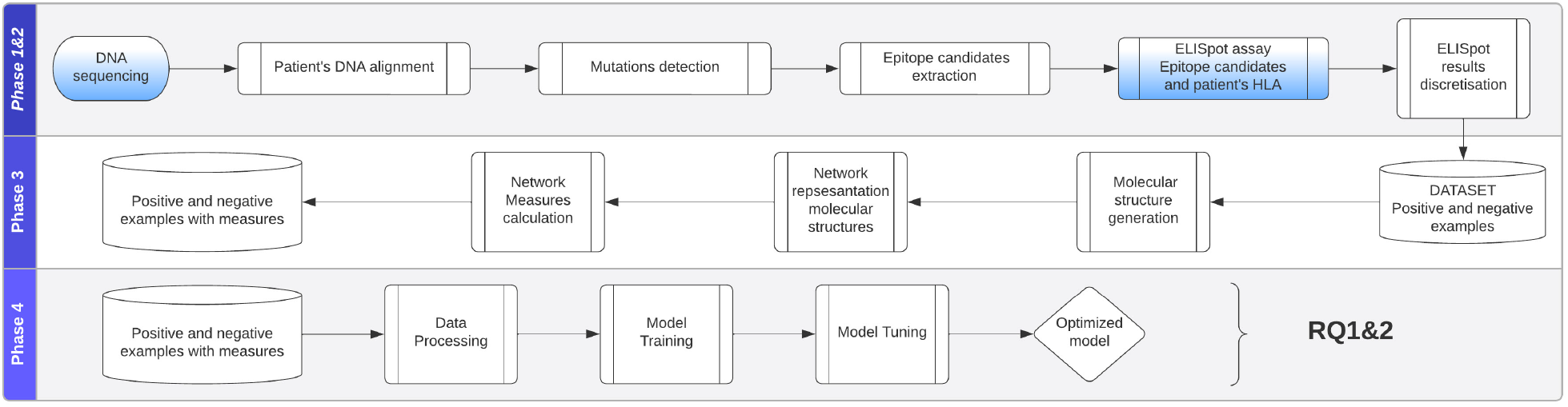
Approach is divided into the following phases: (1) Data Collection, (2) ELISpot assay, (3) Structures and Measurements generation, (4) Analytical approach evaluation

### 3.1 Data

The data, originally published in a study on patient-derived nasopharyngeal cancer (NPC) Organoids for Disease Modeling (28), consists of eighteen NPC tissue samples obtained from patients who underwent biopsy or surgical resection at the National University Hospital Singapore between March 2015 and April 2019. The datasets presented in (28) can be found in online repositories. The names of the repository/repositories and accession number(s) can be found at: NCBI Gene Expression Omnibus.

Specimen collection and experimental use for the study have been approved by the Institutional Review Board of the National Healthcare Group (DSRB Reference: 2015/00098-SRF0004). One part of the tissue collected from patients was immediately transferred to RPMI-1460 media with HEPES and L-Glutamine and 5X antibiotic/antimycotic and 5 µg/ml Metronidazole at 4°C. The other part was fixed in 10% neutral buffered formalin (10% NBF) for routine Hematoxylin and Eosin (HI&E) staining and the remaining tissues were snap-frozen in liquid nitrogen for DNA and RNA extraction. As proof of our methodology, we used mRNA and DNA sequences of 2 patients where one where sequences areas of the mutation have been selected and potential peptide candidates have been identified. Patient one was diagnosed with lung cancer and patient two with nasopharyngeal cancer.

Additionally, enzyme-linked immunospot (ELISpot) assays have been performed with the corresponding patient-specific HLA alleles and peptides producing the quantitative measurements of the immune response (i.e., values we wish to predict).

In order to show the ability of the model to perform well on tumor-specific tasks, we apply the analytical approach to the Cancer Epitope Database and Analysis Resource (CEDAR) (29) containing 1,345,569 Peptidic Epitopes, 116b,026 T Cell Assays, 855,280 B Cell Assays, 4,030,973 MHC Ligand Assays,1,588 Epitope Source Organisms, 652 Restricting MHC Alleles and 4,452 References, originating from cancerrelated studies.

#### 3.1.1 Data overview

The produced data’s attributes as listed in table 1 will give us the ability to create the data structures that represent the relationships between HLA alleles and peptides. This data structure will give us better insights and permit to apply analytical approaches to predict HLA-peptide interactions expressed as a discretized class representing ranges of the numbers of matched blood mononuclear cells (PBMCs).

**Table 1.** Data dictionary of the used data set.

| Attribute | Description |
| --- | --- |
| Peptide Sequence | Represents the structure of amino acids of peptide and is of type string. Every letter represents an amino acid. |
| HLA alleles | The patient's HLA alleles. Represents the HLA protein structure as a sequence of amino acids represented as letters. |
| Matched PBMCs | The number of matched blood mononuclear cells (PBMCs) |
| Discretized PBMC's | PBMC values are discretized into 2 classes: 'no reaction','reaction' |

#### 3.1.2 Generated Molecular Structures

A peptide, as illustrated in figure 5, is a short chain of amino acids (typically 2 to 50) linked by chemical bonds (called peptide bonds). The HLA cell-surface protein, as illustrated in figure 6, is a chain of amino acids that is responsible to regulate the immune system. To understand HLA-peptide interactions we need to understand how the peptide is binding, as illustrated in Figure 7, to the HLA.

**Figure 5.**
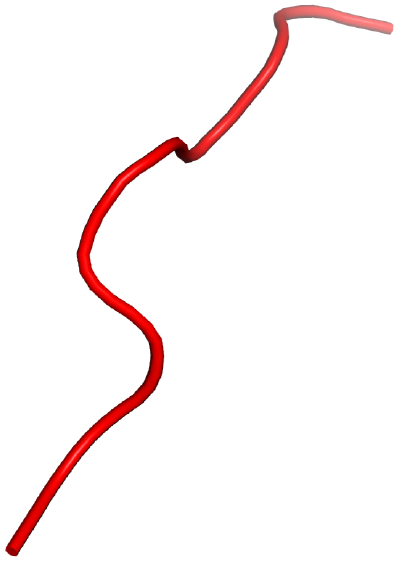
Peptide, a short chain of amino acids, represented in 3-dimensional space.

**Figure 6.**
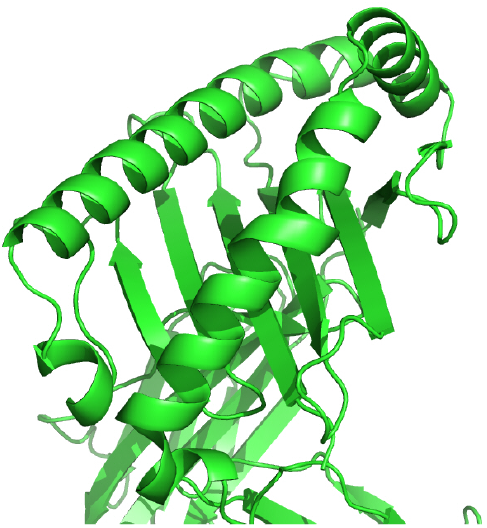
HLA cell-surface protein, chain of amino acids, represented in 3-dimensional space.

**Figure 7.**
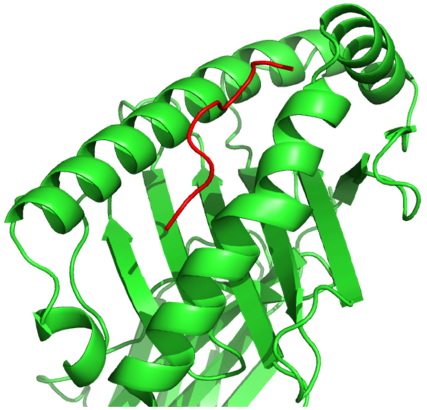
HLA-peptide complex formed by the binding of a peptide to an HLA cell-surface protein, represented in 3-dimensional space.

The HLA-peptide interaction is dependent on the amino acid’s atoms’ chemical reactions that will fall close enough in 3-dimensional space. During phase 2 of our work, as shown in Figure 8, we generate an in-silico representation of the molecular structures composing a system i.e., the patient-specific peptides and HLAs in a system.

**Figure 8.**
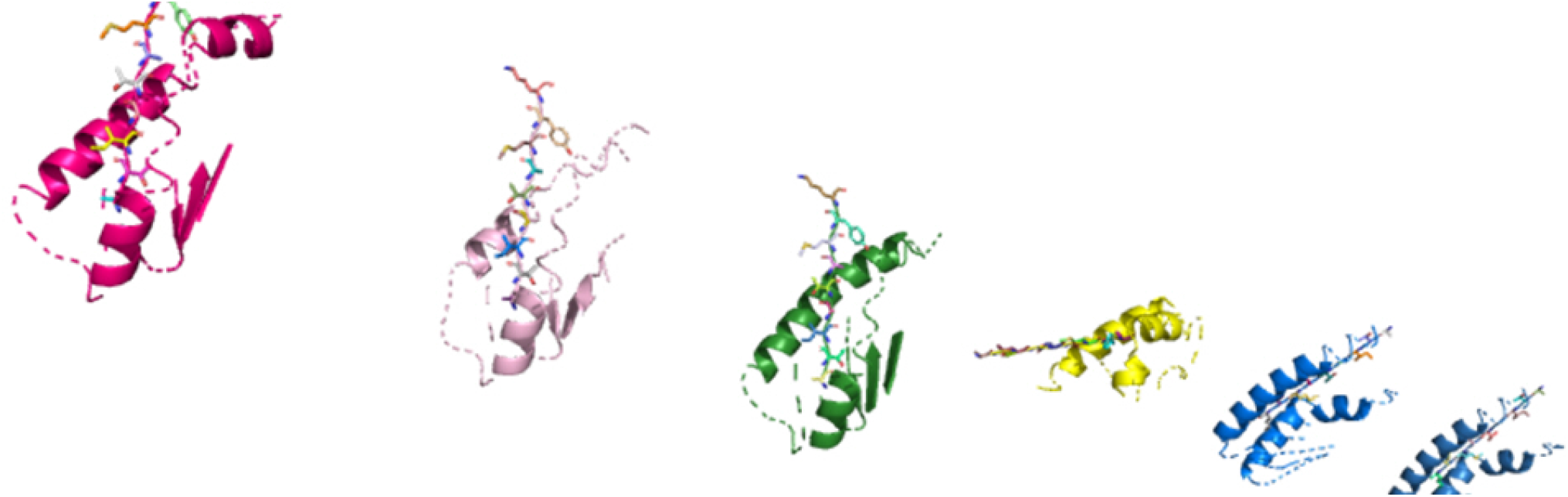
In-silico molecular complexes present in a system, represented in 3-dimensional space.

### 3.2 Generated Network Structures

We chose a network representation with layers to study the diverse relations and interactions between the components. These network representations are called multiplex networks, where a node corresponds to a “physical object,” while node-layer pairs are different instances of the same object.

For instance, a node could represent an online user, while node-layer pairs would represent different accounts of the same user in different online social networks; or a node could represent a social actor, while node-layer pairs would represent different social roles (friend, worker, family member) of the same social actor; or a node could stand for a location in a transportation network, while node-layer pairs would represent stations of different transportation modes (e.g., streets, highways, and subways).

The connection between nodes and node-layer pairs is given by the notion of supra-nodes: i.e., cliques in the supra-graph formed by node-layer pairs that are instances of the same object. To correctly represent a physical object in the different layers of the multiplex network, we break down the peptides into amino acids and the amino acids to their smallest component atoms and their connections bonds. The layers coordinate, atom, monomer, polymer, complex, and system are introduced. The Coordinate layer represents the 3-dimensional coordinates of every atom in the system. The Atom layer, as shown in Figure 9, is the layer that represents the atoms and their bonds that construct objects in the monomer layer, e.g., an amino acid.

**Figure 9.**
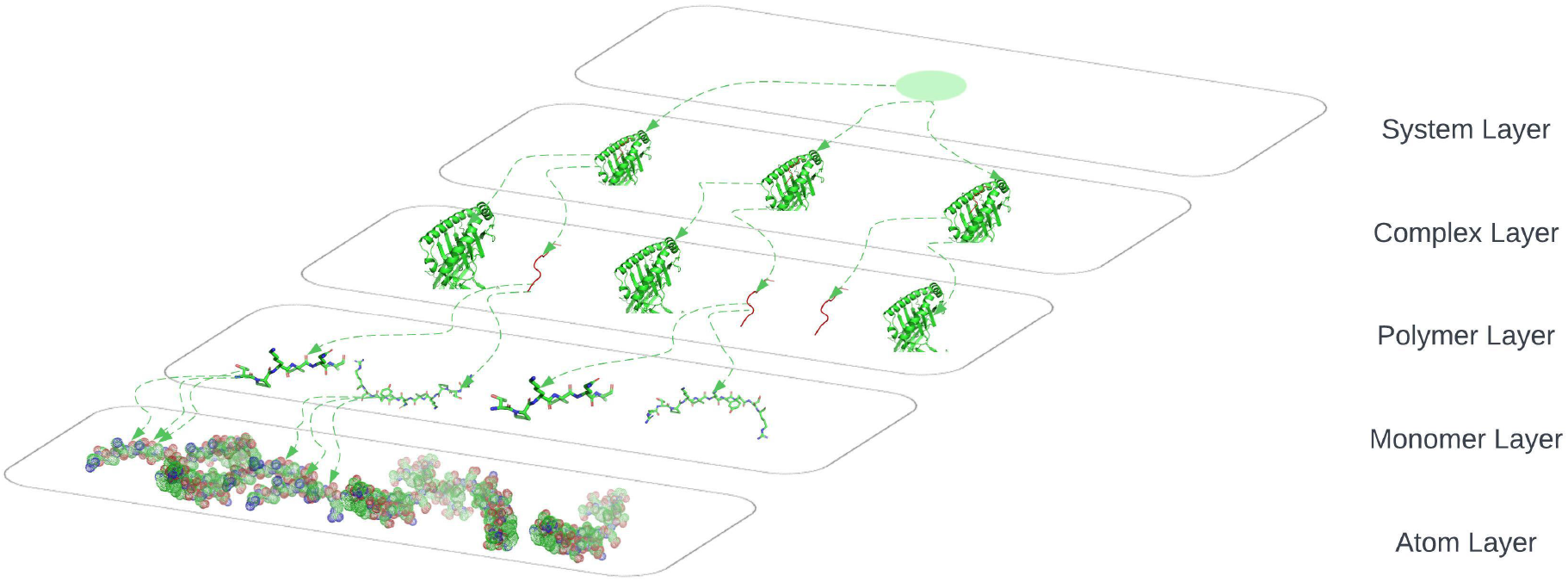
System layer, represents the totality of complexes that exist in an ELISpot assay and is representative of multiple HLA-peptide complexes, represented in 3-dimensional space.

The Monomer layer, as shown in Figure 9, represents the objects of type monomer that are a molecule of any of a class of compounds, mostly organic, that can react with other molecules to form very large molecules or polymers.

The Polymer layer, as shown in Figure 9, represents the objects of type polymer. A polymer is any object of a class of natural or synthetic substances composed of very large molecules, called macromolecules, which are multiples of monomers.

The Complex layer, as shown in Figure 9, represents polymers that form a complex by binding to each other e.g., the peptide binds to an HLA to form a complex as shown in Figure 5.

The System layer represents the totality of complexes that exist in an ELISpot assay and is representative of multiple HLA-peptide complexes. One system could be represented as shown in Figure 9 where we are omitting the coordinates layer.

### 3.3 Networks Measures and Metrics

After generating the graph structures, we calculate network measures on each system. The goal is to select measures such that the analytical model would be able to identify patterns in their values to allow them to distinguish between the two classes of systems i.e., systems that will produce an antibody-peptide epitope and systems that will not produce an antibody-peptide epitope. The used network measures are discussed in detail Appendix Network Measures.

## 4 MODEL TRAINING

### 4.1 Model

We define the prediction of an antibody-peptide epitope as a binary classification problem. Our proposed model takes as input a set of complex networks measures and uses them as features to learn patterns and distinguish between systems that are more likely or less likely to produce an antibody-peptide epitope. Since we seek to discover latent features in the complex networks that represent systems, we choose to rely on deep learning, discussed in Appendix Deep Learning.

Our U-Net with self-attention (UNET-ATT) takes the U-net with attention deep learning architecture previously successfully applied on molecular structures by (30), consisting of blocks of convolutions and deconvolutions, where a convolution block consists of 1 Dimensional Convolutional layer followed by a Max Pooling layer and deconvolution blocks consisting of 1 Dimensional Deconvolutional layer. The visualization of this architecture can be found in Figure 10.

**Figure 10.**
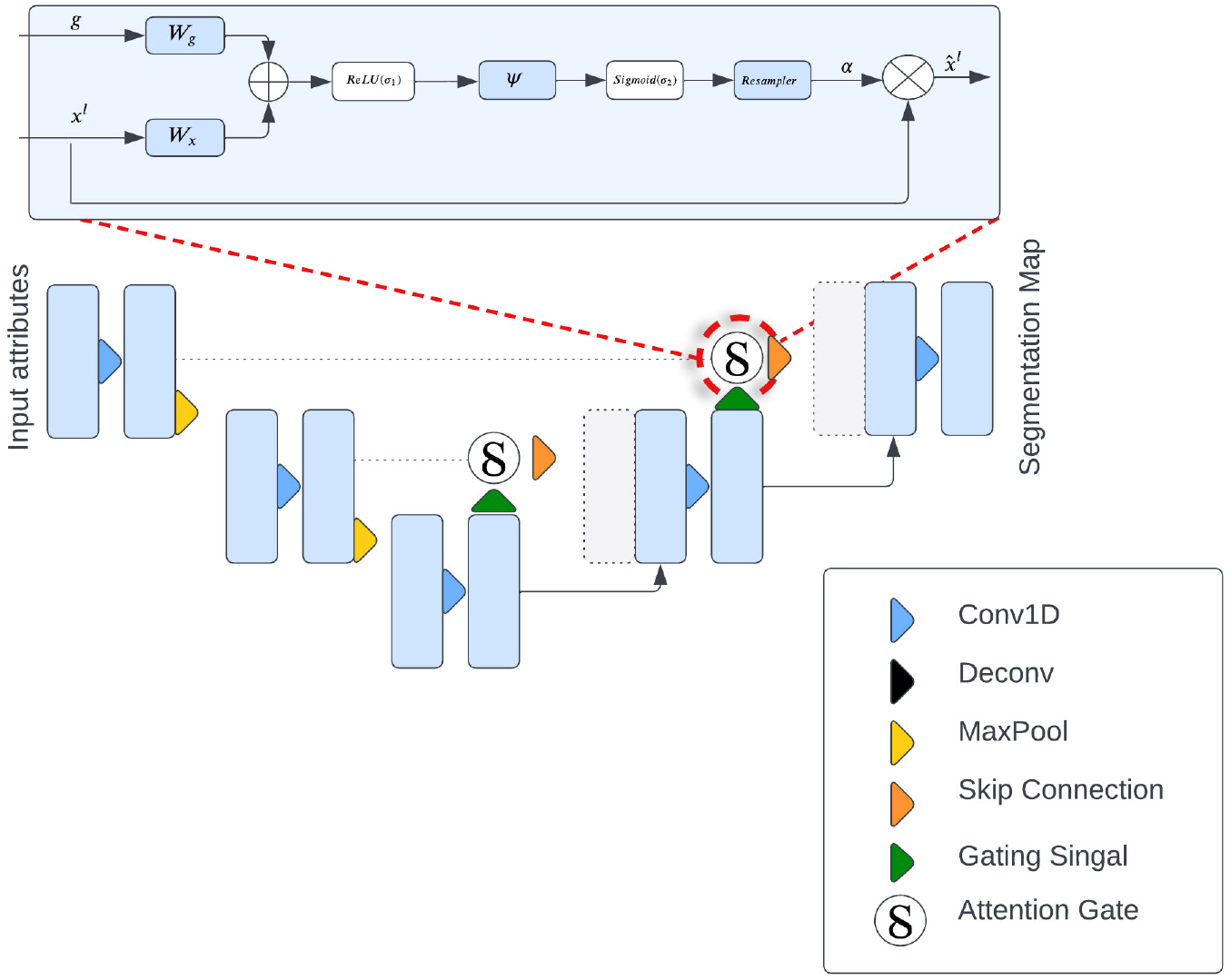
Block diagram of the proposed U-NET with self-attention model architecture. The input feature vector is progressively down-sampled and filtered at each step. Attention gates (AG) filter the latent feature vectors that flow through the skip connections.

The input of the UNET-ATT is a vector with 22 complex networks measures values that undergo batch normalization by introducing additional layers, to stabilize their distribution and control their mean and variance. The batch-normalized inputs flow into a convolution that reduces the input space. Convolutional layers extract higher dimensional representations by processing local features layer-wise. Resulting in the separation of complex network measurements in a high dimensional space based on their semantics. The output value of the layer with input size (*N, C*_*in*_, *L*) and output (*N, C*_*out*_, *L*_*out*_) can be described as:

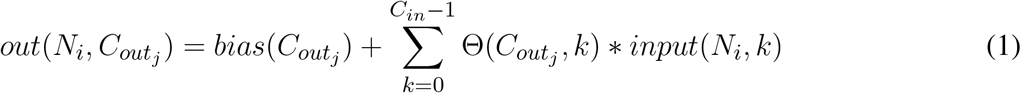

where * is the valid cross-correlation operator, *N* is batch size, *C* denotes the number of channels, *L* is the length of the signal sequence and Θ is the optimizable parameters i.e., weights. The layer output is produced by the sequential application of a linear transformation followed by a non-linear activation function. We apply the Rectified Linear Unit (ReLU) (2) as a non-linear activation function for the convolution layers. To refine the weighting of the latent features, we apply a max-pooling layer.

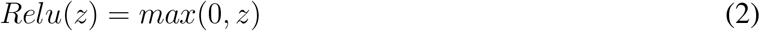

The max pool layer considers the most prominent feature values by eliminating non-prominent features and reducing the feature space. The input to a deconvolution block is the output of the previous deconvolution gated through an attention gate together with the output of a corresponding convolution block. The deconvolution layer performs the opposite transformation to a convolution layer, by applying transformation with the layer parameters to augment the feature space. By minimizing the training objective, e.g., binary cross entropy loss 3, the weights of the layer are adjusted such that it learns to boost important features and makes them more prominent in the network.

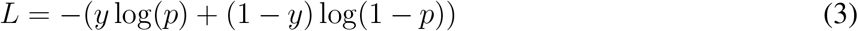

The Deconvolutional network layer applied to the network measures takes as input a feature map *y*^*i*^, composed of *K*_*o*_ feature channels 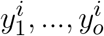. Each channel *c* is represented as linear sum of *K*_*i*_ latent feature maps 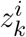 convolved with filter *f*_*k,c*_ :

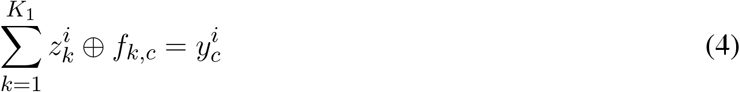

Attention coefficients, *α* ∈ [0.1], are highlighting salient features that are flown through the skip connection see Figure 10. The attention gates output is element-wise multiplication of the input and the attention coefficients: 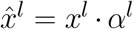 Information extracted from previous layers is transformed through gating to disambiguate irrelevant activations in skip-connections. This is applied before the concatenation in order to merge exclusively important features. Hence, every attention gate optimizes to focus on a set of features, i.e., network measures. As depicted in 10, the gating vector *g* ∈ R is applied on each networks measure in order to identify regions of interest. The attention mechanism is formulated as:

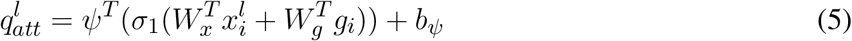

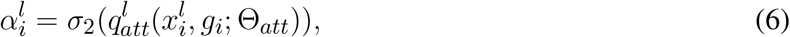

where 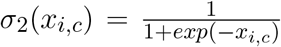 is the sigmoid activation function. The parameters Θ_*att*_ comprised of: linear transformation and bias 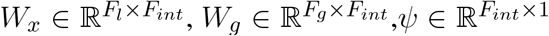 are computed using vectorconcatenation-based attention i.e., channel-wise convolutions of the input vector. The applied randomizing training procedures risk introducing bias into the model by overpredicting classes that are repeatedly fed into the model during training. To avoid this potential overfitting, we introduce a dropout layer that hides a random subset of nodes at each training iteration.

We extend (31) by using the output of the last deconvolution block, a dense representation of non-observable feature maps of the inputted complex networks measures values, as input to the fully connected dense layer in which each node of the layer applies a transformation to output from all nodes from the previous layer. The fully connected layer produces the classification values by producing a probabilistic decision on whether the system will produce an antibody-peptide epitope given the initial inputs i.e., the complex networks measures of the system. The parameters of the UNET-ATT are learnt by minimizing the error, e.g., cross-entropy loss using stochastic gradient descent (SGD).

### 4.2 Data Preparation

To prepare the dataset, we discretize PBMC values and label the ELISpot assays with “1” if the value is above a threshold and is likely to produce an antibody-peptide epitope and “0” if the value is beneath it. Our feature vector consists of the 22 network measure values calculated for the positive and negative examples. For the tumor-specific task, we group the CEDAR data on the tissue type of the Antigen-presenting cells and take epitopes into account with a length between 8 and 25 amino acids, originating from homo sapiens and being part of MHC class I.

## 5 EVALUATION AND DISCUSSION

This section describes our empirical study aimed at evaluating the proposed approach and its main results. We formulated the following research questions:

**Research Question 1**: To what extent is the UNET-ATT model able to correctly predict that a system composed of multiple patient and tumor-specific peptides and HLAs or T cell receptors (TCRs) will result in the presentation of viral epitopes on the surface of the cancer cells in comparison with other models.

**Research Question 2**: To what extent are the models personalized to the patient? Below we discuss the above research questions in detail.

### 5.1 Research Question 1

#### 5.1.1 Approach

To address Research Question 1, we explore the ability of the proposed models to correctly predict if a system composed of multiple patient-specific peptides and HLAs will result in the presentation of viral epitopes on the surface of the cancer cells. Hence, we train individual models for both patients and compare the performance of the Decision tree classifier (DTC), Multi-layer Perceptron classifier (MLPC), Random Forest classifier (RFC), Support Vector Machine (SVM), Naive Bayes classifier (NBC), Logistic Regression classifier (LRC), K-Neighbors Classifier (KNC), U-net with self-attention (UNET-ATT), deep learning on graphs classifier (DLGCL). To evaluate the performance of the algorithms, we use out-of-sample bootstrap validation since this validation technique yields the best balance between the bias and variance compared to single-repetition holdout validation (30).

#### 5.1.2 Results

Four metrics were used to measure the effectiveness of our various models: recall, precision, and F-measure, described in detail in Appendix Evaluation metrics. Precision is the ideal metric to determine the validity of our models and their adequacy in identifying potential systems that will trigger an immune system response. Precision tells the level in which the model was able to correctly identify the systems with high PBMC values in all instances with high PBMC values.

Recall was used, as it allowed us to see the potentially skipped opportunities that the model missed. Utilizing the F-measure statistic, we combined recall and precision into a metric that was able to test the accuracy of the various models. The final metric used to compare the model types was the ROC curve area score, which computes a score from the variability generated by the ROC curve. This metric is useful to show an overall level of model effectiveness, although it summarizes the variability that is shown in the ROC curve. We trained personalized models for patients one and two with their respective data and evaluated them on unseen data. As shown in Table 2, for the personalized models of patient one, the Decision tree classifier, Random Forest classifier, and UNET-ATT performed the best.

**Table 2.** Patient’s one models: Performance of the classifiers for with the highest value in bold.

| Classifier | Precision | Recall | F-measure | PR-AUC |
| --- | --- | --- | --- | --- |
| Decision tree classifier | **1.0** | **1.0** | **1.0** | **1.0** |
| Multi-layer Perceptron classifier | 0.05 | 0.21 | 0.08 | 0.89 |
| Random Forest classifier | **1.0** | **1.0** | **1.0** | **1.0** |
| Support Vector Machine | 0.05 | 0.21 | 0.08 | 0.89 |
| Naive Bayes classifier | 0.88 | 0.86 | 0.83 | 0.92 |
| Logistic Regression classifier | 0.84 | 0.43 | 0.43 | 0.92 |
| K-Neighbors Classifier | 0.88 | 0.86 | 0.83 | 0.92 |
| UNET-ATT | **0.95** | **0.93** | **0.93** | **0.99** |

However, as shown in Table 3, Decision tree classifier and Random Forest classifier did not perform consistently in contrast with UNET-ATT with data from patient two.

**Table 3.** Patient’s two models: Performance of the classifiers with the highest value in bold.

| Classifier | Precision | Recall | F-measure | PR-AUC |
| --- | --- | --- | --- | --- |
| Decision tree classifier | 0.43 | 0.4 | 0.4 | 0.52 |
| Multi-layer Perceptron classifier | 0.05 | 0.21 | 0.08 | **0.89** |
| Random Forest classifier | 0.8 | 0.6 | 0.57 | 0.75 |
| Support Vector Machine | 0.16 | 0.4 | 0.23 | 0.7 |
| Naive Bayes classifier | 0.6 | 0.6 | 0.6 | 0.6 |
| Logistic Regression classifier | 0.43 | 0.4 | 0.4 | 0.52 |
| K-Neighbors Classifier | 0.16 | 0.4 | 0.23 | 0.7 |
| UNET-ATT | **0.87** | **0.8** | **0.8** | **0.83** |

We conjecture that a proper conveyance of the topological attributes, measures, and relationships in complex networks requires sophisticated feature engineering using a neural network classification strategy rather than traditional machine learning classification models. Since the UNET-ATT, consistently outperforms the other models we opt to choose it as our best candidate.

In order to statistically compare the performance of the models, we apply the McNemar test (32) and the Odds Ratio (OR) effect size, where OR larger than 1 indicates that the first technique outperformed the second.

We ran the statistical tests 10 times and compared the results, as shown in Table 4 and Table 5. To accommodate the fact that we performed multiple comparisons, we adjusted the p-values by applying the Bonferonni correction (33).

**Table 4.** Patient One: Statistical comparison between different classification algorithms (McNemar’s test and Odds Ratio). ‘*’ captures the smallest OR among 10 times statistical tests.

| Comparison | p-value | OR |
| --- | --- | --- |
| UNET-ATT vs MLPC | <0.005 | 1.46* |
| UNET-ATT vs RFC | <0.005 | 1.20* |
| UNET-ATT vs SVM | <0.005 | 1.46* |
| UNET-ATT vs NBC | <0.005 | 1.39* |
| UNET-ATT vs LRC | <0.005 | 1.39* |
| UNET-ATT vs KNC | <0.005 | 1.39* |

**Table 5.** Patient Two: Statistical comparison between different classification algorithms (McNemar’s test and Odds Ratio). ‘*’ captures the smallest OR among 10 times statistical tests.

| Comparison | p-value | OR |
| --- | --- | --- |
| UNET-ATT vs MLPC | <0.005 | 1.37* |
| UNET-ATT vs RFC | <0.005 | 1.37* |
| UNET-ATT vs SVM | <0.005 | 0.92* |
| UNET-ATT vs NBC | <0.005 | 1.36* |
| UNET-ATT vs LRC | <0.005 | 0.91* |
| UNET-ATT vs KNC | <0.005 | 0.91* |

In this context, we formulate our null hypothesis for each test as there is no statistically significant difference between the performance of the two algorithms and the alternative hypothesis as there is a statistically significant difference between the performance of the two algorithms. We adjusted the alpha value to 0.005 to perform the hypothesis test at a 5% significance level.

As we can observe in Table 4 and Table 5, the McNemar test results show that the null hypothesis is rejected as there are statistically significant differences (*p <* 0.5*/*10) in the performance of the UNET-ATT model compared to all the other models. The OR values show that the UNET-ATT model has more chances to correctly predict that a system will result in the presentation of viral epitopes on the surface of the cancer cells from the other methods.

We further apply the analytical approach to the CEDAR dataset, which contains 1,345,569 tumor-related peptidic epitopes, where we create tumor-specific UNET-ATT models by grouping the data based on the cell tissue type of the antigen-presenting cells.

In Table 6, we present the performance metrics of the tumor-specific UNET-ATT models trained on antigen-presenting cell tissue type-specific data. The UNET-ATT model can generalize with acceptable F-measure performance on all tissue types, when trained on specific tissue types it archives remarkable F-measures for Breast and Lung and is acceptable for Lymphoid.

**Table 6.** Tumor-specific classifiers: Performance of the classifier on tumor-specific tasks.

| Tumor specific classifier | Accuracy | Precision | Recall | F-measure | PR-AUC |
| --- | --- | --- | --- | --- | --- |
| All tissue types | 0.69 | 0.66 | 0.76 | 0.70 | 0.69 |
| Blood | 0.56 | 0.60 | 0.47 | 0.52 | 0.56 |
| Breast | 1 | 1 | 1 | 1 | 1 |
| Lung | 0.90 | 1 | 0.86 | 0.92 | 0.92 |
| Lymphoid | 0.71 | 0.67 | 0.76 | 0.71 | 0.72 |

### 5.2 Research Question 2 Personalized model

#### 5.2.1 Approach

To address Research Question 2, we apply statistical analysis techniques such as Multivariate analysis of variance (MANOVA) on the network measures to evaluate if they are statistically significantly different per patient. Additionally, we look at how different network measures correlate with PMBC values per patient. We apply a structure learning algorithm to the data to learn the structure of the directed acyclic graph (DAG) to analyze the causality of data features, i.e., network measures and the matched PBMC values. Finally, we compare the performance of personalized models on unseen data from different patients to understand if models can generalize over unseen data from different patients. The applied statistical analysis techniques are discussed in detail in Appendix Statistical Analysis.

#### 5.2.2 Results

We can conclude that data attributes have a patient-specific distribution and correlation from the performed data exploration on the network measures. This finding agrees with the general understanding that tumor mutations and immune system reactions are patient and tumor-specific and that a personalized approach is required. Additionally, the data exploration shows that network attributes of the molecular structures are patient and tumor-specific and that some show correlations with statistical significance with the produced PMBC values. That indicates that a machine learning approach that relies on these attributes as input can be applied to produce predictions personalized towards a patient. Finally, we compared the performance of personalized models on unseen data from different patients. We observed that they do not generalize well to data from different patients and perform better on unseen data from the same patient.

## 6 THREATS TO VALIDITY

### 6.1 Internal and Construct Validity

Our analysis is mostly threatened by the generation of the molecular structures in a system, as the wrong placement of molecular structures in a complex may result in different network measure values. However, not all network measures will be impacted, and the “skewing” of these network measure values will be consistent over all the data systems. As for the selection of a threshold for the PBMC values such that they are split into positive and negative systems i.e., systems that produce an antibody-peptide epitope and systems that don’t, this threshold can be modified by domain experts without resulting in a drastic change of model performance given that the data is well balanced and sampled. The same reasoning can be applied to choosing to predict more than two classes i.e., splitting the PBMC values into buckets and predicting a range of PMBC values within which a system would fall.

### 6.2 External Validity

Our analysis was performed on data from 2 patients with the same tumor. However, we cannot claim the generality of our observations to other tumors or all patients with the concrete tumor of our data. Further investigation is needed on data from multiple tumors and more patients to mitigate this threat.

## 7 CONCLUSION

Predicting the specific antibody peptide that will be presented on the surface of a tumor cell is of paramount importance. It permits the creation of personalized treatments that will trigger the human immune system to destroy the tumor cell. In this study, we proposed a novel data structure leveraging multiplex networks and derive network measures as attributes leveraging the theory and methods of complex networks, together with a deep learning approach for optimal feature engineering and personalized antibody-peptide epitope binary classification. Our results reveal that machine learning and deep learning models are able to binary classify antibody-peptide epitopes based on the derived attributes from the proposed data structure.

In particular, the proposed UNET-ATT demonstrates an F-measure of 0.8 and 0.93 for personalized models for patients one and two, respectively. Additionally, the UNET-ATT model consistently outperforms the other models on both patients in contrast with machine learning baseline models that show different results for different patients. Additionally, UNETT-ATT can generalize on large data set comprising all tissue types with acceptable F-measures. The same approach demonstrated F-measures of 0.92 and 1.0 for Lung and Breast antigen-presenting tissue types, respectively, proving its ability to specialize in tumor-specific tasks.

Additionally, we analyzed the validity of building personalized models and found that data attributes have patient-specific distribution and correlation. The data exploration shows that network measures of the molecular structures are patient and tumor-specific and that some show a correlation with statistical significance with PMBC values. Finally, we compared the performance of models personalized toward a patient on unseen data from different patients. We observed that they do not perform well in contrast to their performance on unseen data from the same patient.

These findings agree with the general understanding that tumor mutations and reactions of the immune system are patient and tumor-specific and that a personalized approach is required for optimal results. However, we need to caution that these findings cannot be generalized to other tumors or even other patients with the same tumor from our data, given that the sample size is too small. Further investigation is needed on data from multiple tumors and more patients to mitigate this threat.

## 8 APPENDIX

## 8.1 Network Measures

## 8.1.1 Attribute Assortativity Coefficient Source

Assortativity measures the similarity of connections in the graph with respect to the attribute source (34). In the generated multiplex structure, the attribute “source” refers to the unique id of the node the edge originates from.

### 8.1.2 Degree Pearson Correlation Coefficient

One measure of the degree assortativity of a network is the Pearson correlation coefficient (35) between degrees across edges *r*. Similar to the Pearson correlation coefficient, it varies between −1 ≤ *r* ≤ 1: for *r <* 0, the network is assortative, for *r* = 0 the network is neutral and for *r* > 0 the network is disassortative.

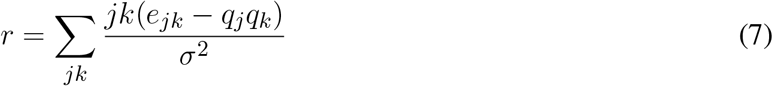

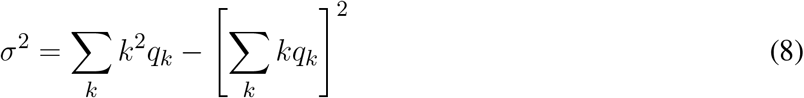

#### 8.1.3 Closeness Centrality

The closeness centrality of a node *u* is the reciprocal of the average shortest path distance to *u* overall *n* − 1 reachable nodes (34).

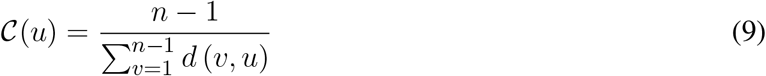

where *d*(*v, u*) is the shortest-path distance between *v* and *u*, and *n* is the number of nodes that can reach *u*. Concretely we average the betweenness centrality.

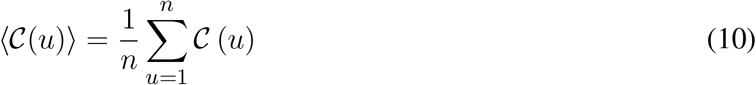

#### 8.1.4 Betweenness Centrality

Betweenness Centrality (34) measures the importance of a node based on the number of paths between two other nodes that pass through it. Betweenness Centrality is a guide to the influence nodes have over the flow of information, and energy between others. Betweenness Centrality *x*_*i*_ is given by

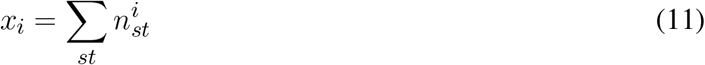

where 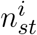 is the node *i* that lies on the shortest path from vertices *s* to *t*. Concretely we average the betweenness centrality

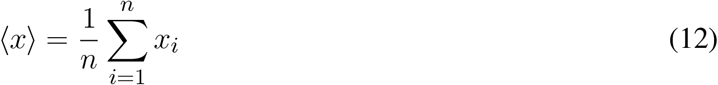

#### 8.1.5 Density

The density of a graph is the ratio between the edges present in a graph and the maximum number of edges that the graph can contain. Conceptually, it provides an idea of how dense a graph is in terms of edge connectivity.

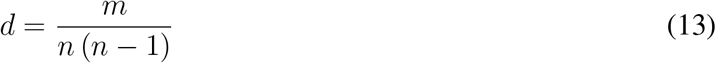

where *n* is the number of nodes and *m* is the number of edges in a graph *G*.

#### 8.1.6 Average Core Hubs

Average core hubs is a measure of the average of the degrees of the top 100 hubs in a system: where *k* is the degree of vertice *i*.

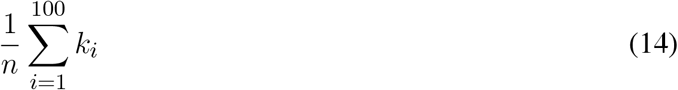

#### 8.1.7 Average Degree

The average degree of a network:

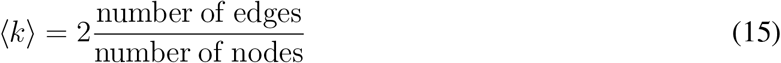

is related to the density of the network, wherein in a dense network the average degree grows linearly with the number of nodes, while in a sparse network, it grows sublinearly.

#### 8.1.8 Transitivity

The fundamental type of relation between nodes in a network is “connected by an edge.” If the “connected by an edge” relation were transitive it would mean that if node *u* is connected to node *v*, and *v* is connected to *w*, then *u* is also connected to *w*.

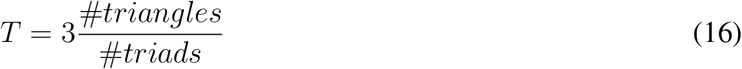

where “triads” are two edges with a shared vertex and triangles are loops of length three.

#### 8.1.9 Global Clustering Coefficient

The clustering coefficient (34) is a measure of the degree to which vertices in a network tend to create tightly knit groups, closed triads:

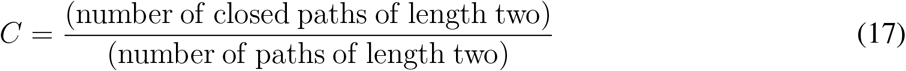

*C* = 1 implies perfect transitivity, a network whose components are all closed triads. *C* = 0 implies no closed triads. Concretely we apply the Watts and Strogatz defined clustering coefficient that quantifies the likelihood that two nodes that are connected to the same node are also connected to each other. the degree to which vertices in a network tend to create tightly knit groups, closed triads:

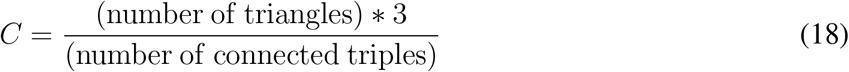

Concretely we apply the following equation

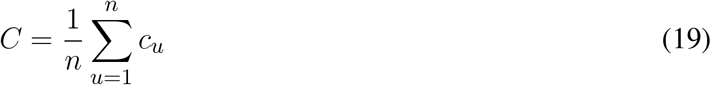

where *n* is the number of nodes and *c*_*u*_ is:

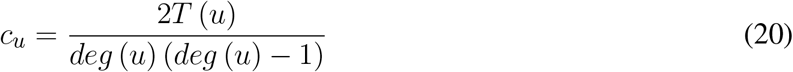

where *T* (*u*) is the number of triangles through node *u* and *deg* (*u*) is the degree of *u*.

#### 8.1.10 Size largest Component

In undirected networks, typically exists, large components that fill most of the network, while the rest of the network is divided into a lot of small, disconnected components. (35) The size of the largest connected component can be expressed by:

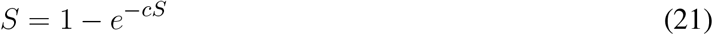

introduced by Erdos and Renyi in 1958, where *c* denotes the mean degree or the average of in- and outgoing edges. We compute the size of the largest component S of each system and look at its distribution and correlation with the particular patient and the measured matched PBMC i.e., ELISpot values.

#### 8.1.11 Average Spectral clustering

A fast approach to separate a network in communities is by applying spectral modularity maximization by assigning a node to one of two groups of communities. If we consider s as a vector in n-dimensional space, that implies that the vector must point to one of the corners of the n-dimensional hypercube. By creating a relaxed method where *s* is allowed to take any value of 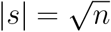 the form we arrive at a matrix notation *Bs* = *βs*. Hence the optimal *s* is one of the eigenvectors of the modularity matrix and *β* is the corresponding eigenvalue. We can find which eigenvector by applying

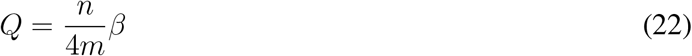

Since *s*^*T*^ *s* = *n* we can maximize the inner product *s*^*T*^ *u* = Σ_*i*_ *s*_*i*_*u*_*i*_. The maximum is achieved when *s*_*i*_*u*_*i*_is positive for all *i*, which occurs when *s*_*i*_ has the same sign as *u*_*i*_ for all *i*:

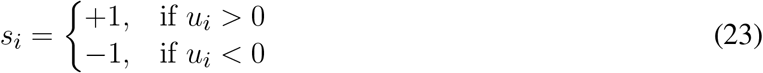

This leads to a simple approach; we calculate the eigenvector of ton the modularity matrix corresponding to the highest eigenvalue and assign nodes to communities according to the signs of the elements in this vector. We slightly modify it by assigning zero to the nodes that correspond to negative eigenvalues instead of -1:

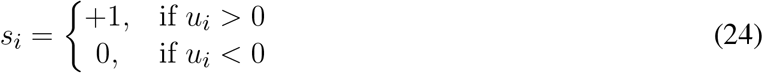

Then we calculate the average 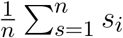of the assigned values that produces a value between zero and one and is indicative of how balanced the distribution of the nodes between two communities is.

#### 8.1.12 Mean distance

If we consider an undirected network with *d*_*ij*_ as the length of the shortest path through the network between nodes *i* and *j* or otherwise called distance between *i* and *j*. Then the mean distance *l*_*i*_ between nodes *i* and *j* is:

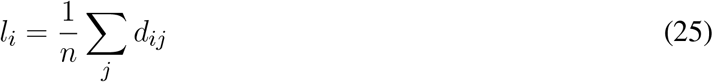

Then the mean distance *l* between nodes for the whole network is the average of this quantity over all nodes

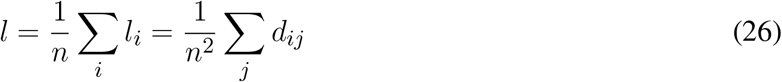

#### 8.1.13 Degree Distribution

The degree distribution is defined as the fraction of the nodes that have a particular degree *k*. Following the frequency probability approach, we can say that the probability of k or 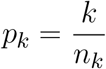 is the number of nodes with degree k. We can use *p*_*k*_, for example, to find the number of nodes with such a degree with *np*_*k*_.

It turns out that many real-world networks have degree distributions with a tail of high-degree hubs like this. In the language of statistics, we say that the degree distribution is right-skewed. One such distribution is the power law distribution: *p*_*k*_ = *Ak*^*−α*^, where *α* and *A* are constants, which are used for modeling scale-free networks.

#### 8.1.14 Cumulative Distribution of Degree

A way of properly visualizing a power-low distribution is to construct the cumulative distribution function, which is defined as

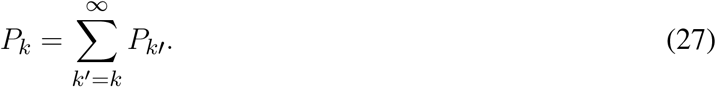

Or *P*_*k*_ is the fraction of nodes that have degree *k* or greater or alternatively as probability*P*_*k*_ that a randomly chosen node has a degree *k* or greater.

#### 8.1.15 Walks, loops, and paths

A walk is any sequence of nodes such that every consecutive pair of nodes in the sequence is connected by an edge. In general, a walk can intersect itself, revisiting nodes it has visited before. Walks that do not intersect themselves are called paths or self-avoiding walks. The length r of a walk in a network is the number of edges (not nodes) traversed along the walk. The number of walks of a given length r in a network can be generalized to: Laplacian clustering

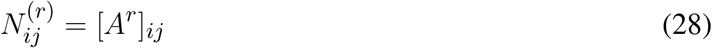

where […]_*ij*_ denotes the *ij*^*th*^ element of the matrix and *A* is the adjacency matrix.

Loops is a special type of walk that start and end at the same node i. The total number of loops of length r is the sum of this quantity over all possible starting points i:

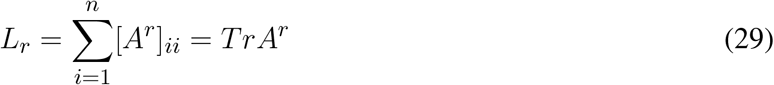

where *Tr* stands for trace 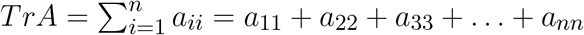

#### 8.1.16 Paths lengths

A graph diameter is the longest and shortest path in a graph. The diameter of the graph varies with the n number of nodes as ln n. (natural log of n). The basic idea of the calculation of the diameter is that if we grow a set of nodes in a graph by iteratively adding neighbors to the set, the number of neighbors grows up by a factor of *c*, the average degree 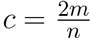; where *m* is the number of edges in the graph and n is the number of nodes, on average per iteration *s*.

So, the average number of nodes one step away is *c*, and by growing by a factor of *c* every iteration then the average number of nodes *s* iterations away is *c*^*s*^. Given the exponential growth, it does not take long until the number of nodes reached is equal to the total number of nodes in the graph *c*^*s*^ ≃ *n* or equivalently 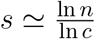 implying that the diameter is approximately 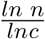

#### 8.1.17 Degree centrality

A large volume of work is dedicated to centrality. The question is which are the most important vertices in or central nodes in a network. There are many possible definitions of importance and consequently many centrality measures. Degree, the number of incoming and outgoing edges a node has, is sometimes called degree centrality to emphasize its use as a centrality measure. Useful though it is quite a crude measure as it awards a node one centrality point for every neighbor it has. But not all neighbors are necessarily equivalent.

#### 8.1.18 Eigenvector centrality

In many circumstances, a node’s importance in a network is increased by having connections with other nodes that are themselves important. Eigenvector centrality is an extension of degree centrality that takes this factor into account. Eigenvector centrality awards to a node several points proportional to the centrality scores of its neighbors. The eigenvector centrality *x*_*i*_ of node *i* is defined to be proportional to the sum of the centralities of i’s neighbors

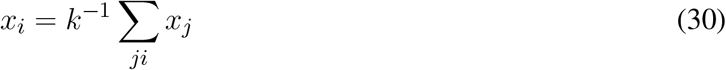

An alternative way of writing using the adjacency matrix is:

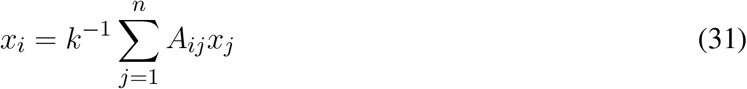

In matrix notation

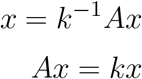

where *x* is the vector with elements equal to the centrality scores *x*_*i*_ and *k* is a constant. In other words, *x* is an eigenvector of the adjacency matrix.

#### 8.1.19 Katz centrality

One solution is to grant every node with a small amount of centrality, regardless of its position:

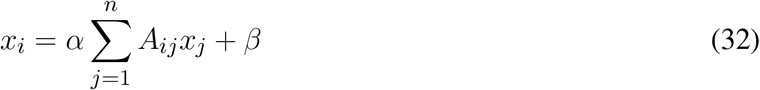

where *α, β* are positive constants.

#### 8.1.20 Page Rank Centrality

If a node with high Katz centrality has edges pointing to many others, then all of those others also get high centrality. A high-centrality node pointing to one million others gives all one million of them high centrality. One could argue that this is not always appropriate. In many cases, it means less if a node is only one among many that are pointed to. The centrality gained by virtue of receiving an edge from a prestigious node is diluted by being shared with so many others. We can derive the following variant of Katz centrality in which the centrality we derive from our network neighbors is proportional to their centrality divided by their out-degree.

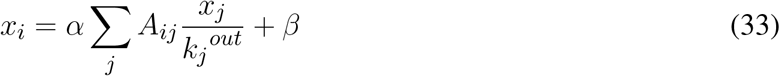

#### 8.1.21 Hubs and authorities

The concept of hubs and authorities in networks was first put forward by Kleinberg and developed by him into a centrality algorithm called hyperlink-induced topic search or HITS. The HITS algorithm gives each node *i* in a directed network two different centrality scores, the authority centrality *x*_*i*_ and the hub centrality *y*_*i*_, which quantify nodes’ prominence in the two roles.

The defining characteristic of a node with high authority centrality is that it is pointed to by many nodes with high hub centrality. Conversely, the defining characteristic of a node with high hub centrality is that it points to many nodes with high authority centrality. In Kleinberg’s approach the authority centrality of a node is defined to be proportional to the sum of the hub centralities of the nodes that point to it:

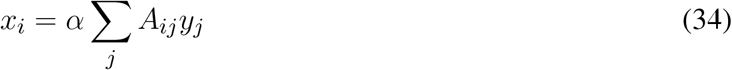

where *α* is a constant. Similarly, the hub centrality of a node is proportional to the sum of the authority centralities of the nodes it points to:

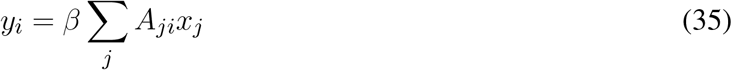

Thus, the authority and hub centralities are respectively given by eigenvectors of *AA*^*T*^ and *A*^*T*^ *A* with the same eigenvalue.

#### 8.1.22 Multiplex Participation Coefficient

As we are operating in the context of a Multiplex data structure, it is necessary to introduce measurements that can capture and quantify the heterogeneity and the richness of the connectivity patterns between layers. A suitable measure to compute is the distribution of the degree of a node among the various layers. This value will be equal to zero if all the node’s links are in the same layer, on the other hand, it will take its maximum value if the edges are uniformly distributed over the layers.

The multiplex participation coefficient (36) follows the same principle to quantify the participation of a node in different layers of the network. By measuring values in [0, 1] it quantifies whether the edges to a vertex are uniformly distributed amongst all the layers.

Concretely, the coefficient *P*_*i*_ is equal to zero when all the edges of *i* are in the same layer, while *P*_*i*_ = 1 when vertex *i* has the same number of edges on each of the M layers. The participation coefficient of the multiplex is defined as the average of the participation coefficients of all the vertices, i.e.,

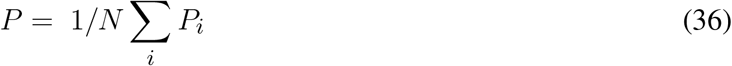

#### 8.1.23 Multiplex interdependence

To capture the multiplex contribution to the reachability of each layer of the network (36), the node interdependence is introduced as:

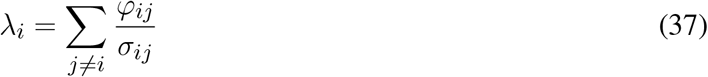

where *σ*_*ij*_ is the total number of shortest paths between node *i* and node *j* on the multiplex network. *φ*_*ij*_ is the number of shortest paths between node *i* and node *j* that use links in two or more layers. Hence, the node independence is 1 when the shortest paths make use of edges in at least two layers, and 0 when each of the shortest paths uses only the layers in the system. Averaging*λ*_*i*_ over all nodes, produces the multiplex interdependence, i.e.,

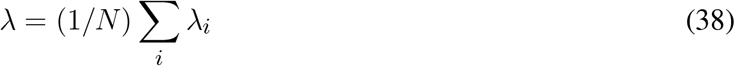

#### 8.1.24 Supracentrality

Introduced by (37) Supracentrality is a generalization of eigenvector-based centralities, centrality measures that include PageRank, hub and authority scores, and eigenvector centrality. Coupling centrality measures across layers with weighed interlayer edges create a Supracentrality matrix *C* (*w*), where w controls the extent to which centrality changes over the layers.

### 8.2 Machine Learning Algorithms

#### 8.2.1 Cluster Analysis K-means

Cluster analysis divides data into groups (clusters) that are meaningful, useful or both. If meaningful groups are the goal, then the clusters should capture the natural structure of the data. In clustering analysis, there are multiple types of techniques to identify and group objects together. The Prototype-based clustering techniques create a one-level partitioning of the data objects where a cluster is a set of objects in which each object is closer (more similar) to the prototype that defines the cluster than to the prototype of any other cluster. For data with continuous attributes, the prototype of a cluster is often a centroid, i.e., the average (mean) of all the points in the cluster. The K-means clustering technique is simple, and we begin with a description of the basic algorithm. We first choose K initial centroids, where K is a user-specified parameter, namely, the number of clusters desired. Each point is then assigned to the closest centroid, and each collection of points assigned to a centroid is a cluster. The centroid of each cluster is then updated based on the points assigned to the cluster. We repeat the assignment and update steps until no point changes clusters, or equivalently until the centroids remain the same.

#### 8.2.2 Cluster Analysis K-means

For the purpose of classification, we are interested in computing the probability of observing a class label y for a data instance given its set of attribute values x. This can be represented as *P* (*y*|*x*), which is known as the posterior probability of the target class. The Naïve Bayes classifier assumes that the class-conditional probability of all attributes x can be factored as a product of class-conditional probabilities of every attribute *x*_*i*_, as described in the following equation:

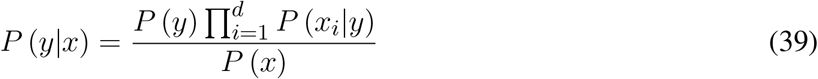

Since *P* (*x*) is fixed for every y and only acts as a normalizing constant to ensure that *P* (*y x*) in [0, 1], we can write

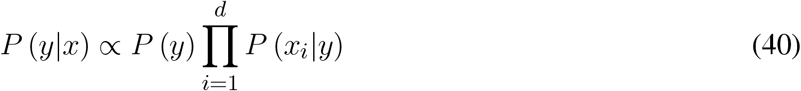

Hence, we are looking for a class *c* ∈ [*Y, N*] that maximizes 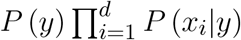 where *x*is an instancefrom the dataset with d attributes. To use the classification approach, we utilize the discrete classes as described in Table 1 as ground truth.

#### 8.2.3 Decision tree classifier

To solve a classification problem, we can ask a series of carefully crafted questions about the attributes of the instance. Each time we receive an answer, we could ask a follow-up question until we can conclusively decide on its class label. The series of questions and their possible answers can be organized into a hierarchical structure called a decision tree. The tree has three types of nodes: A root node, with no incoming links and zero or more outgoing links. Internal nodes, each of which has exactly one incoming link and two or more outgoing links. Leaf or terminal nodes, each of which has exactly one incoming link and no outgoing links. Every leaf node in the decision tree is associated with a class label. The non-terminal nodes, which include the root and internal nodes, contain attribute test conditions that are typically defined using a single attribute. Each possible outcome of the attribute test condition is associated with exactly one child of this node.

#### 8.2.4 Deep Learning

A multi-layer neural network generalizes the basic concept of a perceptron to more complex architectures of nodes that are capable of learning nonlinear decision boundaries. A generic architecture of a multi-layer neural network where the nodes are arranged in groups called layers. These layers are commonly organized in the form of a chain such that every layer operates on the outputs of its preceding layer. In this way, the layers represent different levels of abstraction that are applied to the input features in a sequential manner. The composition of these abstractions generates the final output at the last layer, which is used for making predictions.

During the training process of deep learning models, we iterate over mini-batches and minimize the loss for the mini-batch. We can empirically verify that the loss is going down while the accuracy is going up per iteration step. Several challenges arise during the training and selection of a model. The biggest is the bias-variance trade. We refer to bias as the error we introduce when we try to approximate a real-life task with a mathematical function i.e., a machine learning or deep learning model. When the model can successfully approximate a task, it becomes more flexible and fits well the data as such bias reduces. Variance refers to the amount the error changes when the model is presented with new data. Ideally, we wish that the difference in performance on different data sets is minimal and that the model generalizes well on the task. Usually, as the models become more flexible, the variance will increase, and the bias will decrease. In other words, the models will be able to approximate the task better but will not generalize on unseen data. As we train the models, the bias tends to initially decrease faster than the variance increases. Consequently, the expected test error declines. However, at some point, increasing flexibility has a negligible impact on the bias but starts to significantly increase the variance. When this happens the test error increases. We say that the model is overfitting on the training data, it has learned it by heart, and will not generalize well.

Especially large deep learning models can suffer from this phenomenon as the substantial number of parameters provides for the ability to become more flexible to the training data set. For example, in the field of Computer Vision, it is common that a model starts to wrongly associate a specific background with a class of an object and will misclassify objects based on their background.

### 8.3 Statistical Analysis

#### 8.3.1 ELISpot value Distribution

In Figure 11, we analyze the distribution of the ELISpot values for both patients. The medians from both patients are different as well as the distribution of the values. The values of patient one are restricted to a smaller range and show an outlier in the upper quartile. In contrast patient two shows values distributed over a wider range, again with a wider distribution in the upper quartile. We can conclude that the patients show different distributions of observed PBMC values, i.e., the PBMC values are patient-specific.

**Figure 11.**
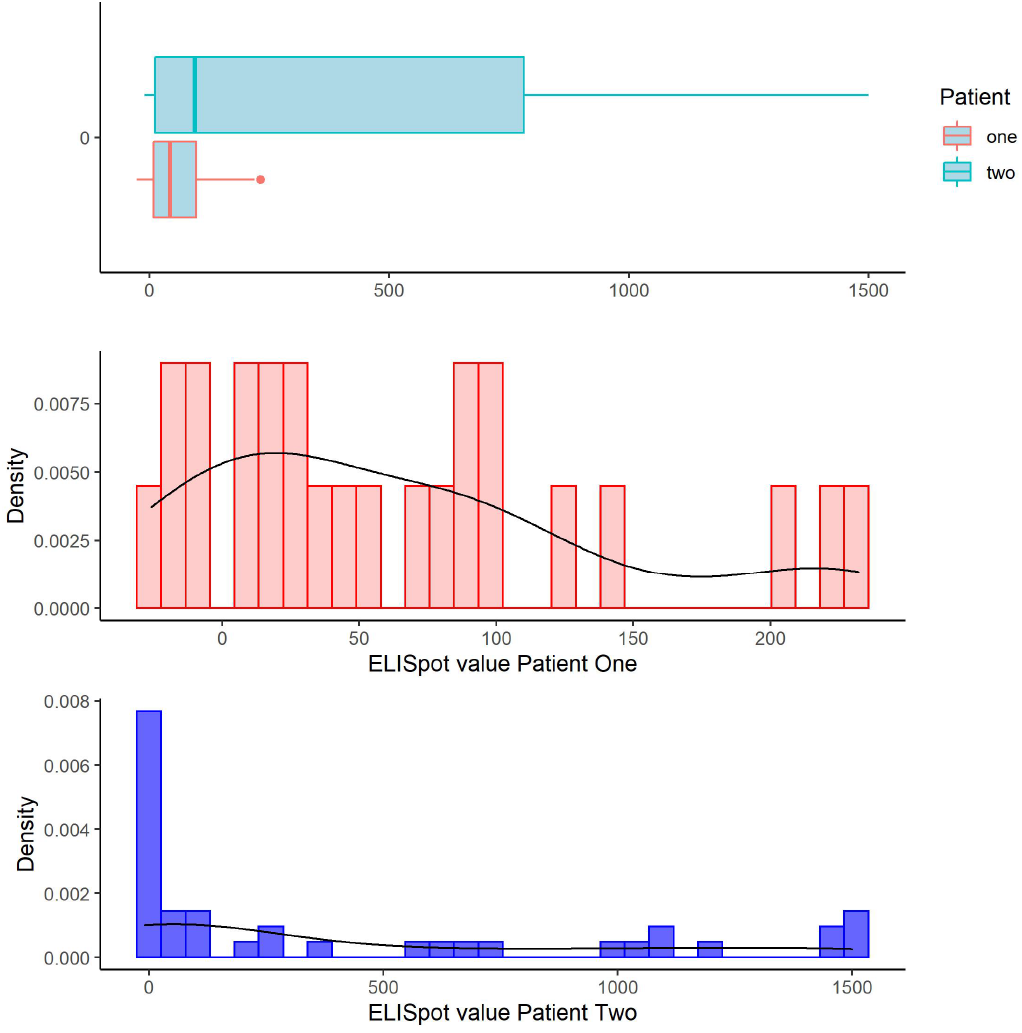
ELISpot histogram plot, depicting ELISpot values for both patients. The medians from both patients are different as well as the distribution of the values.

#### 8.3.2 Peptides number versus ELISpot Value

In Figure 12, we graph a violin plot to analyze how the number of peptides in a system correlates with the PBMC values and specific patients. We see that patient two has systems with exclusively one peptide in contrast to patient one which has systems with one, two, and three peptides. Patient one has a high concentration of systems with only one peptide. The correlation between the number of peptides and ELISPot values is not clear. Systems with one peptide seem to produce higher PBMC values in contrast to systems with multiple peptides. This conclusion should be taken with caution since only patient one has systems with more than one peptide.

**Figure 12.**
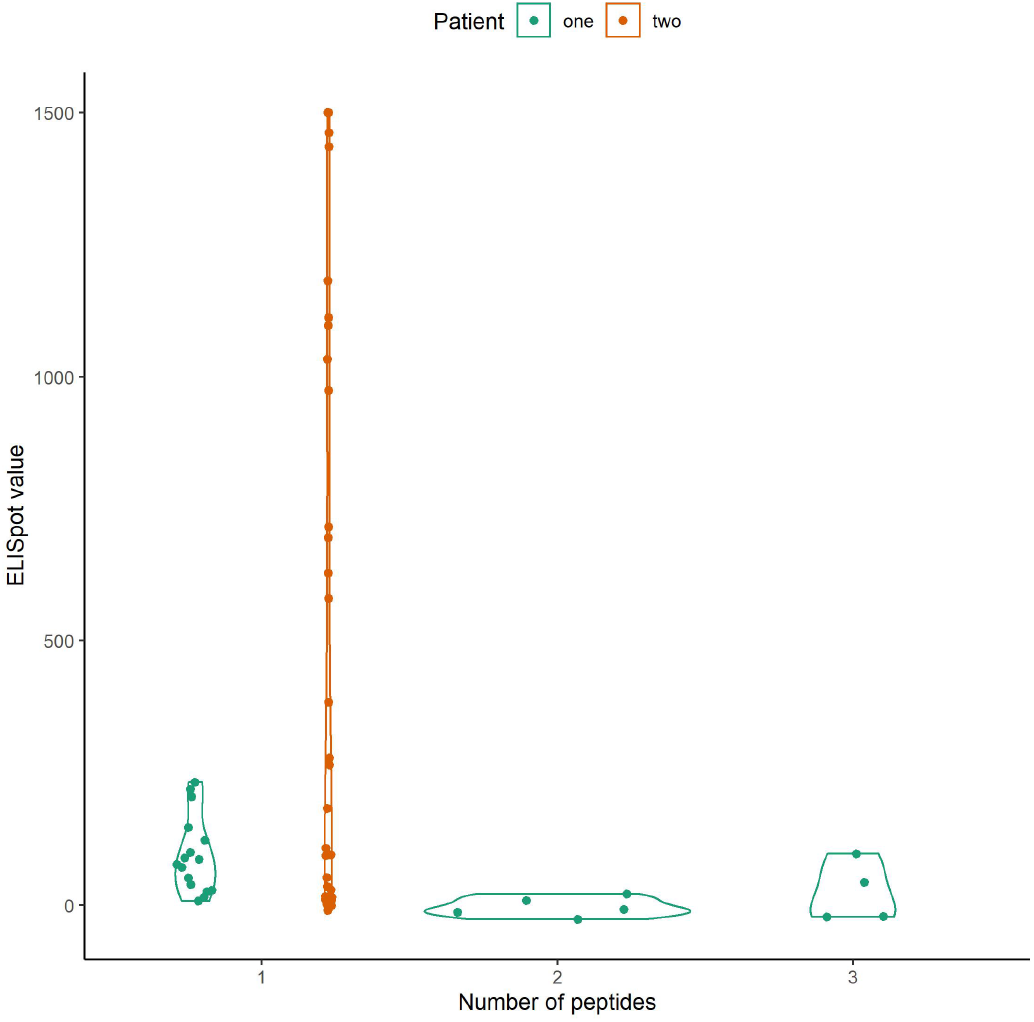
ELISpot violin plot, depicting ELISpot values for both patients. In order to analyze how the number of peptides in a system correlates with the PBMC values and specific patients.

#### 8.3.3 Number of Amino acids versus ELISpot values

In Figure 17, we analyze the distribution of ELISpot values in relation to the total number of Amino Acids originating from the peptides in a system. We look at these correlations for both patients individually and together. Here the difference between the patients is not so clear, we observe a pattern in both patients where the lower and the upper range of the number of patients correlate with a high ELISPot value. It is noticeable that systems with 30 amino acids consistently produce PBMC values above 500 for patient two. We can conclude that systems with the higher total number of amino acids in the peptides tend to produce higher ELISpot values.

#### 8.3.4 Linear Model coefficient analysis

We apply a linear regression model to better understand the potential influence of particular data attributes. We can understand the predictor’s influence on the produced PBMC values by analyzing the coefficients. From the results shown in Table 7, we can see that the total number of amino acids in a system has a positive linear relationship with the produced PBMCs for patient one and negative for patient two. The model’s performance differs between the two patients, and the same coefficients have a different linear relationship on the produced PBMC for the two patients. Hence the relationship between these coefficients and the matched PBMC value is patient-specific. This observation indicates that tumor mutations and immune system response are not only tumor but also patient-specific.

**Figure 17.**
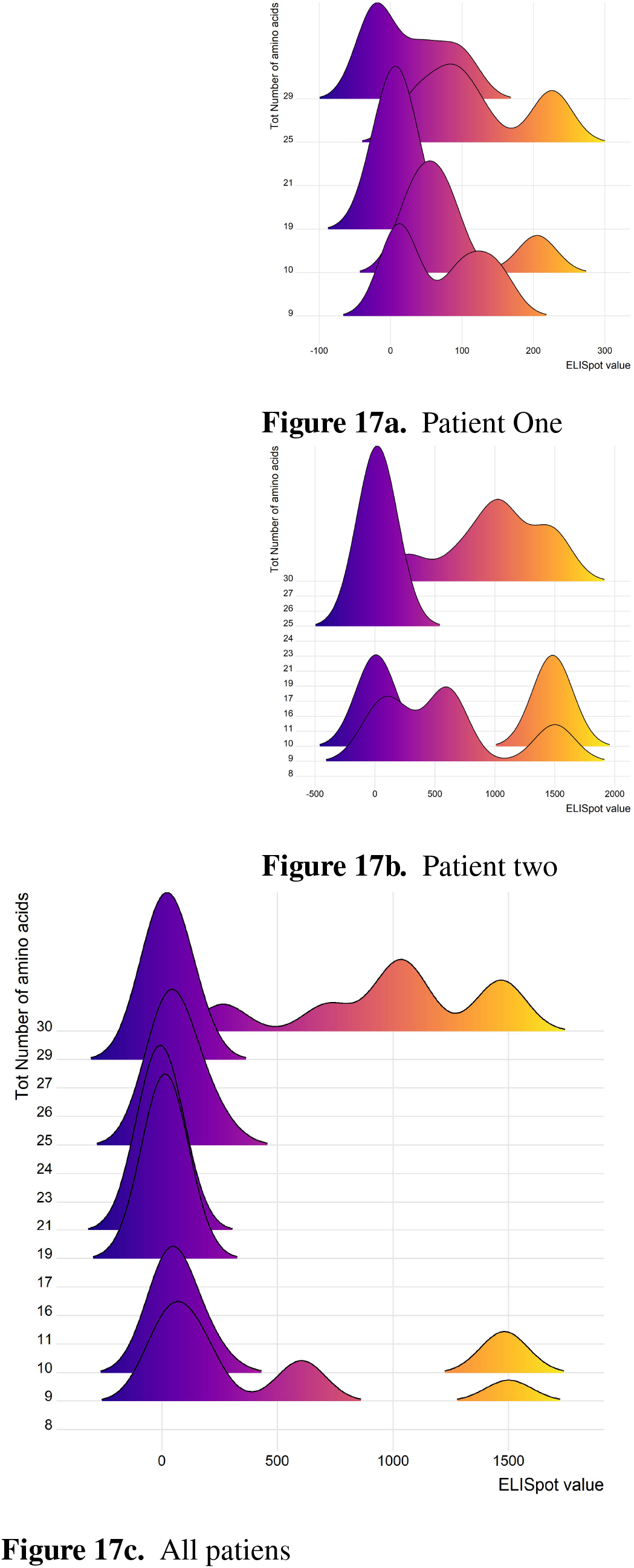
Distribution of ELISpot values in relation to the total number of Amino Acids originating from the peptides in a system. **(A)** Distribution of Patient one ELISpot values in relation to the total number of Amino Acids. **(B)** Distribution of Patient two ELISpot values in relation to the total number of Amino Acids.**(C)** Distribution of both patients’ ELISpot values in relation to the total number of Amino Acids.

**Table 7.**
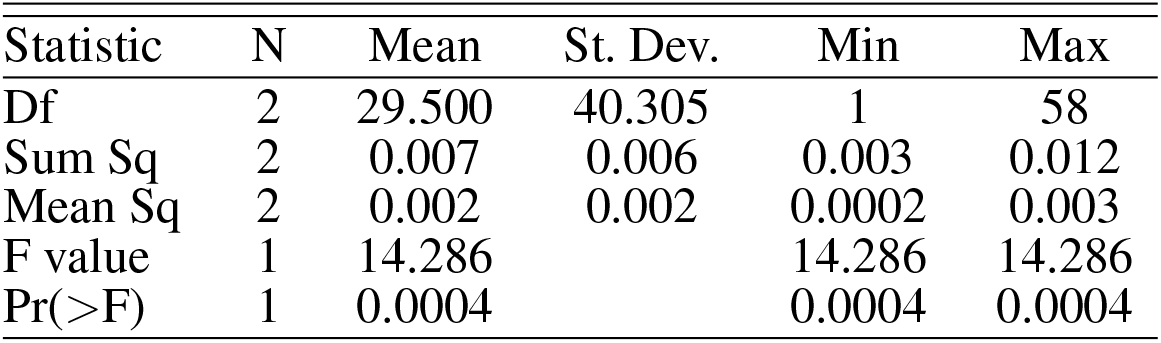
Response Size largest Component

| Statistic | N | Mean | St. Dev. | Min | Max |
| --- | --- | --- | --- | --- | --- |
| Df | 2 | 29.500 | 40.305 | 1 | 58 |
| Sum Sq | 2 | 0.007 | 0.006 | 0.003 | 0.012 |
| Mean Sq | 2 | 0.002 | 0.002 | 0.0002 | 0.003 |
| F value | 1 | 14.286 |  | 14.286 | 14.286 |
| Pr(>F) | 1 | 0.0004 |  | 0.0004 | 0.0004 |

#### 8.3.5 Multivariate analysis of variance

Multivariate analysis of variance (MANOVA) is a statistical procedure for the comparison of multivariate sample means. It is often used when there are two or more dependent variables, and we wish to perform regression analysis and analysis of variance for them by one or more factor variables or covariates. In our case, we wish to determine which network measures are highly significantly different among patients. From the results shown in Tables 8 to 14, we see that Matched PBMC, Average Core hubs, and Average Betweenness Centrality have a high statistical significance with a p-value < 0.001. Closeness Centrality, Degree Pearson Correlation Coefficient, Page Rank Centrality, Attribute Assortativity Coefficient Source with p-value < 0.01 Size largest Component, Response Degree Centrality, Density statistical significance with p-value < 0.05.

**Table 8.** Response Closeness Centrality

| Statistic | N | Mean | St. Dev. | Min | Max |
| --- | --- | --- | --- | --- | --- |
| Df | 2 | 29.500 | 40.305 | 1 | 58 |
| Sum Sq | 2 | 0.00000 | 0.00000 | 0.00000 | 0.00000 |
| Mean Sq | 2 | 0.00000 | 0.00000 | 0.00000 | 0.00000 |
| F value | 1 | 10.164 |  | 10.164 | 10.164 |
| Pr(>F) | 1 | 0.002 |  | 0.002 | 0.002 |

**Table 9.**
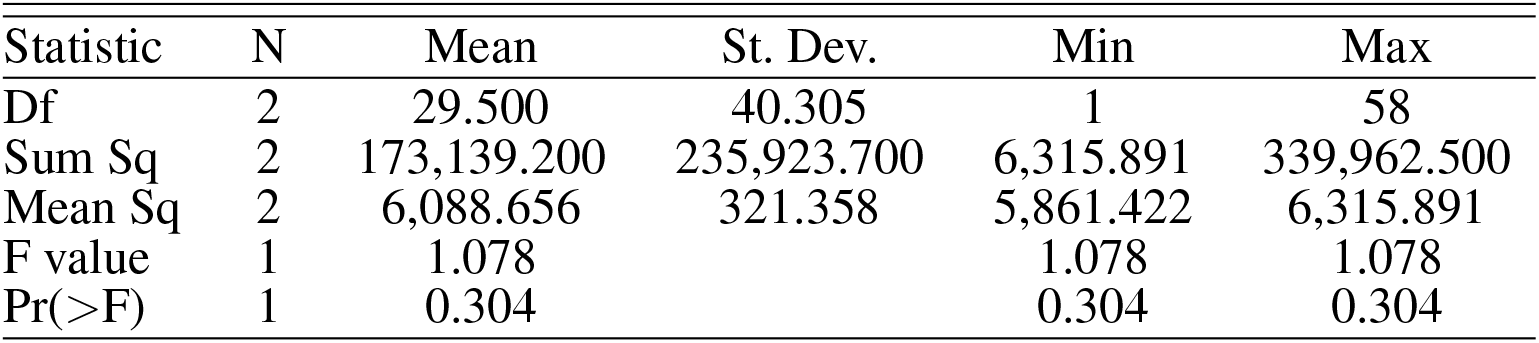
Response Degree Pearson Correlation Coefficient

**Table 10.** Response Eigenvector Centrality

| Statistic | N | Mean | St. Dev. | Min | Max |
| --- | --- | --- | --- | --- | --- |
| Df | 2 | 29.500 | 40.305 | 1 | 58 |
| Sum Sq | 2 | 0.00000 | 0.00000 | 0.00000 | 0.00000 |
| Mean Sq | 2 | 0.000 | 0.000 | 0.000 | 0.00000 |
| F value | 1 | 13.680 |  | 13.680 | 13.680 |
| Pr(>F) | 1 | 0.0005 |  | 0.0005 | 0.0005 |

**Table 11.** Response Page Rank Centrality

| Statistic | N | Mean | St. Dev. | Min | Max |
| --- | --- | --- | --- | --- | --- |
| Df | 2 | 29.500 | 40.305 | 1 | 58 |
| Sum Sq | 2 | 0.003 | 0.003 | 0.001 | 0.005 |
| Mean Sq | 2 | 0.001 | 0.001 | 0.0001 | 0.001 |
| F value | 1 | 11.914 |  | 11.914 | 11.914 |
| Pr(>F) | 1 | 0.001 |  | 0.001 | 0.001 |

**Table 12.** Response Attribute Assortativity Coefficient Type

| Statistic | N | Mean | St. Dev. | Min | Max |
| --- | --- | --- | --- | --- | --- |
| Df | 2 | 29.500 | 40.305 | 1 | 58 |
| Sum Sq | 2 | 0.002 | 0.002 | 0.001 | 0.004 |
| Mean Sq | 2 | 0.0003 | 0.0003 | 0.0001 | 0.001 |
| F value | 1 | 7.848 |  | 7.848 | 7.848 |
| Pr(>F) | 1 | 0.007 |  | 0.007 | 0.007 |

**Table 13.** Response Attribute Assortativity Coefficient Source

| Statistic | N | Mean | St. Dev. | Min | Max |
| --- | --- | --- | --- | --- | --- |
| Df | 2 | 29.500 | 40.305 | 1 | 58 |
| Sum Sq | 2 | 0.0001 | 0.0001 | 0.00000 | 0.0001 |
| Mean Sq | 2 | 0.00000 | 0.00000 | 0.00000 | 0.00000 |
| F value | 1 | 0.046 |  | 0.046 | 0.046 |
| Pr(>F) | 1 | 0.830 |  | 0.830 | 0.830 |

**Table 14.**
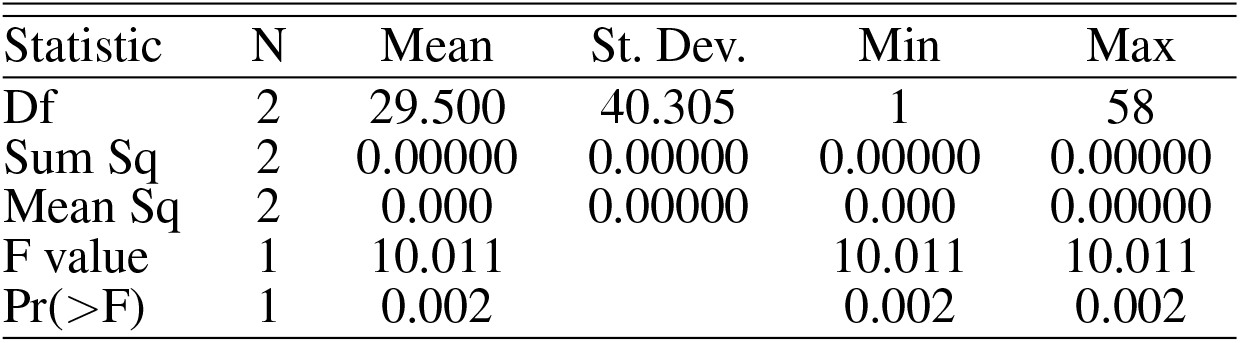
Response Matched PBMC

#### 8.3.6 Size largest Component

As visualized in Figure 18, even though the medians of both patients are close, the values of patient one are better distributed with a large portion in the lower quartile. Patient two shows a high concentration around the median and the upper quartile and an outlier in the upper whisker.

**Figure 18.**
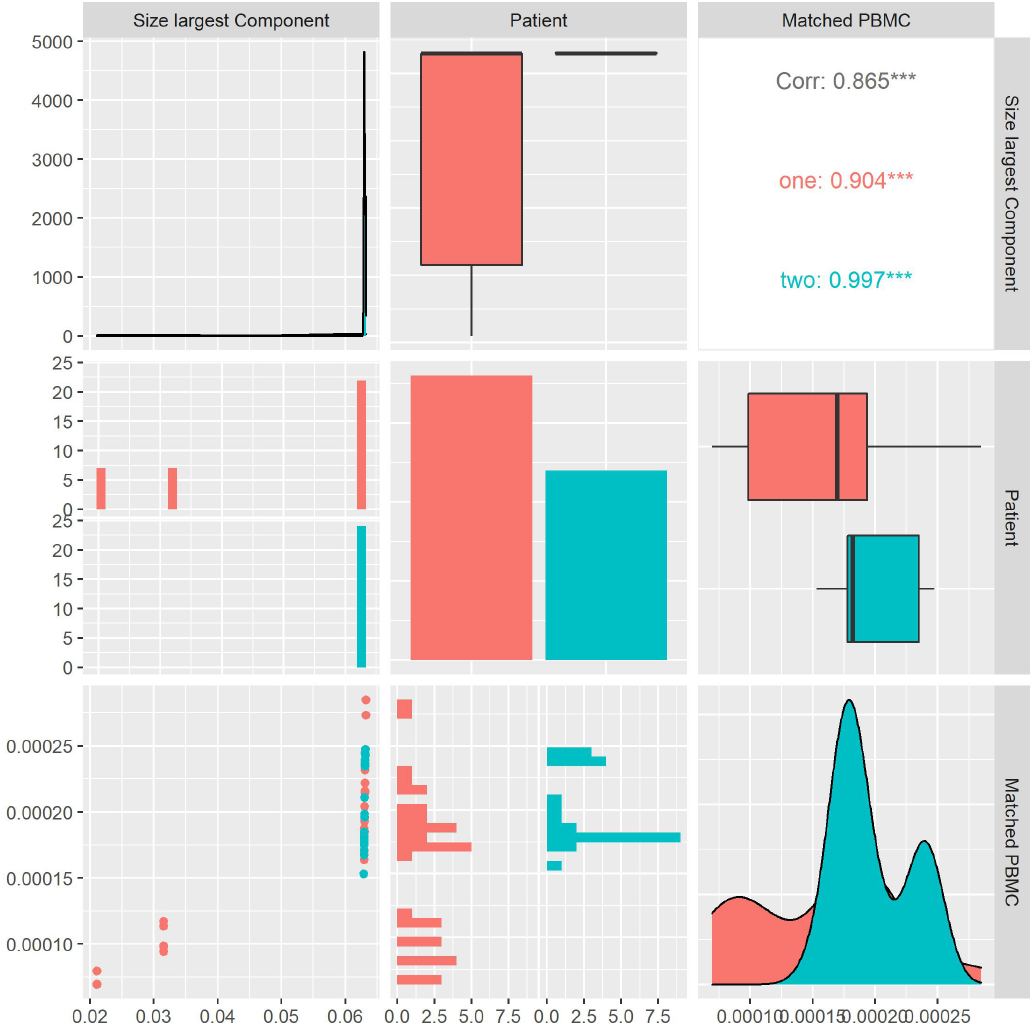
Patient Matched PBMC’s vs. Size largest Component.

When considering the S values over all the systems, we observe a positive correlation with the matched PBMCs with statistical significance shown by a p-value¡0.10. This could mean that systems that create larger connected components have a higher chance of producing a higher PBMC value. This is confirmed by the distribution of the PBMC values of patient two, showing overall significantly higher values than patient one, and the distribution of the size of the largest components that is concentrated around the median of 30 and the upper quartile. When looking at the individual patients, we observe that patient two shows an inverse correlation, even though not significant, with the matched PBMCs. Patient one shows a positive correlation with matched PBMCs.

#### 8.3.7 Average Spectral clustering

As visualized in Figure 19, we calculate the average spectral clustering for each system and look at its distribution and correlation with the patient and the measured matched PBMC, i.e., ELISpot values. We see that the distributions for both patients are quite similar. Patient two produces values skewed toward the lower quartile, while patient one values are better distributed around the median. Even though the combined data of both patients show an inverse Pearson correlation coefficient with the matched PBMCs, patient one shows a positive correlation with statistical significance indicated by a p-value < 0.05. This might indicate a difference in the produced PMBCs between patients with similar avg. spectral clustering values. In other words, the systems of individual patients react differently to a similar internal node distribution between two communities.

**Figure 19.**
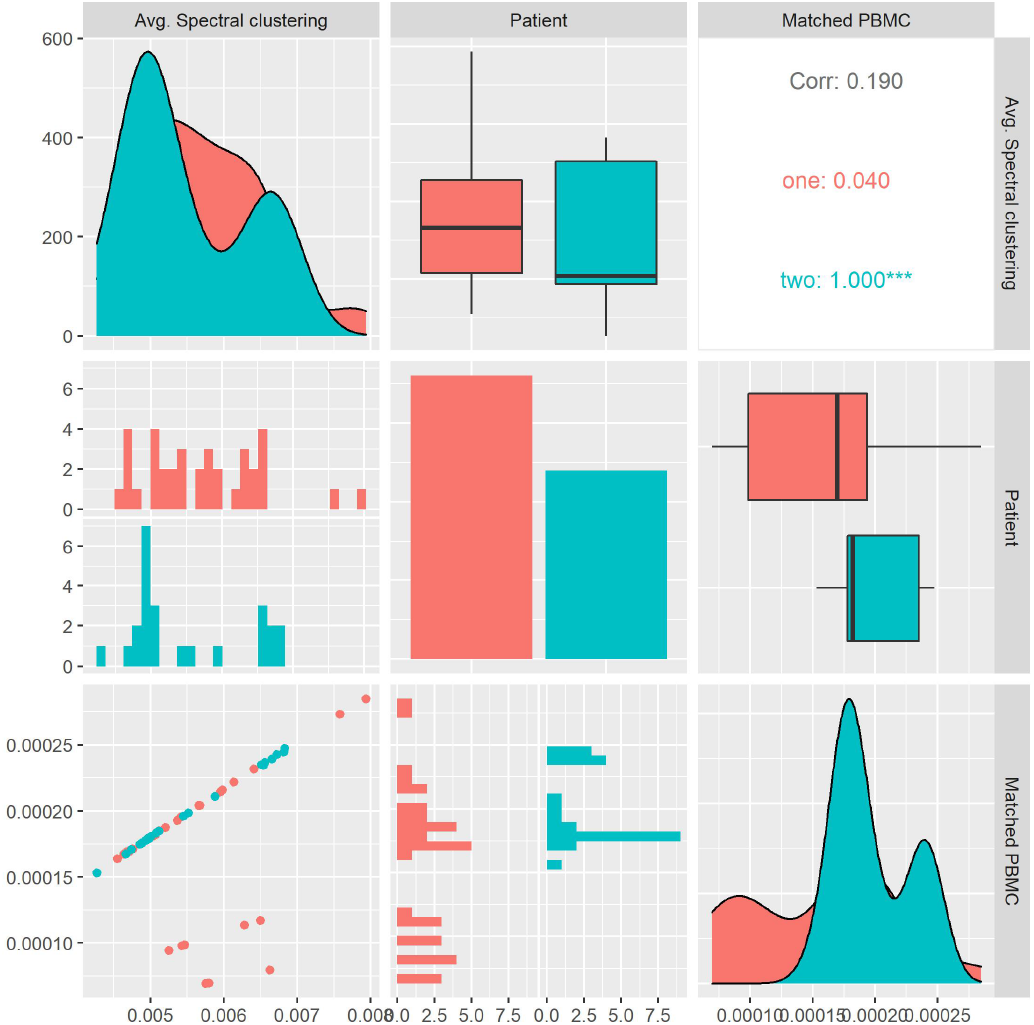
Patient Matched PBMC’s vs. Avg. Spectral clustering

#### 8.3.8 Global Clustering Coefficient

As visualized in Figure 20, we calculate the global clustering coefficient for each system and look at its distribution and correlation with the particular patient and the measured matched PBMC i.e., ELISpot values. We see that the distributions for both patients are quite similar. Patient two produces values that are skewed toward the upper quartile while patient one values are better distributed around the median.

**Figure 20.**
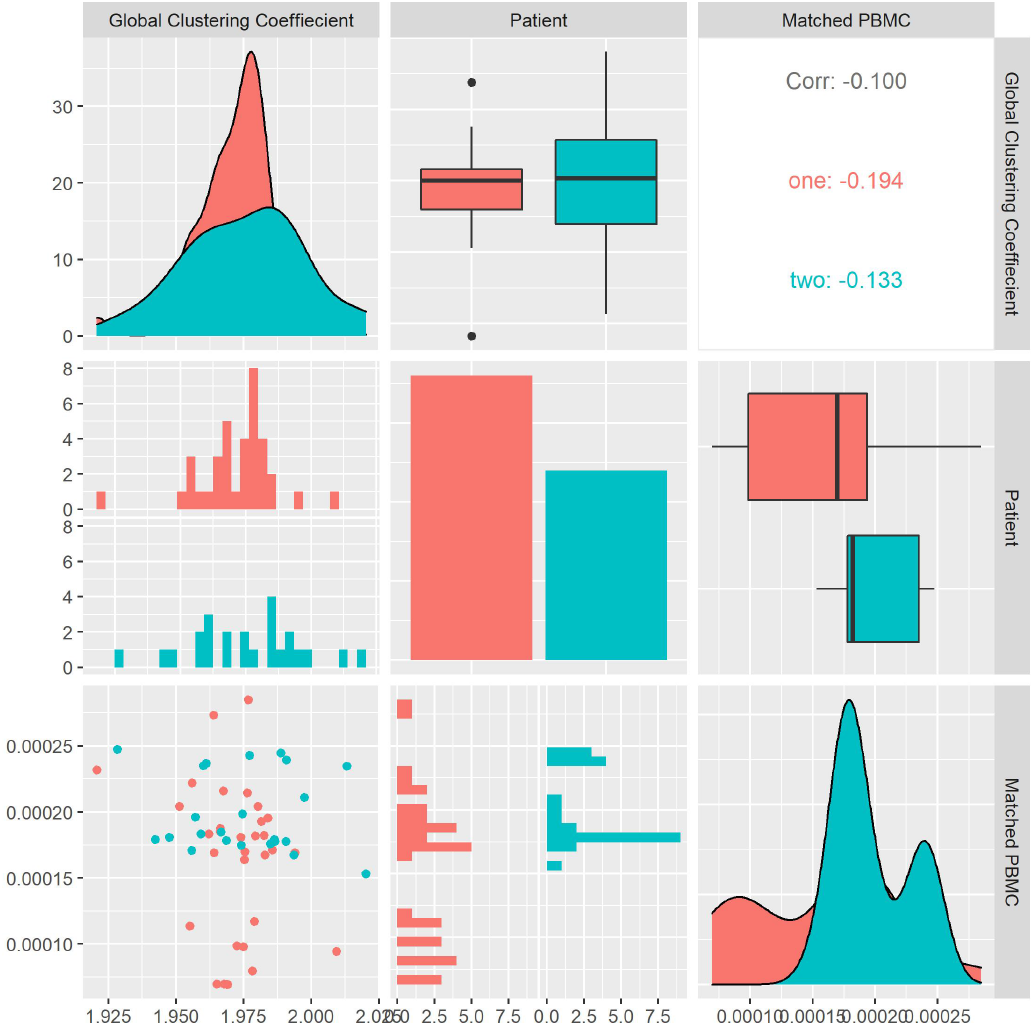
Patient Matched PBMCs vs. Global Clustering Coefficient

Combined data of both patients show a positive Pearson correlation coefficient, with statistical significance indicated by a p-value < 0.05, with the matched PBMCs. Patients one and two show a similar positive correlation. This observation can imply that systems with high clustering coefficients, i.e., systems where monomers, polymers, and atoms that are connected to the same monomer, polymer, or atom have a high likelihood to be connected to each other, will produce a higher number of matched PBMCs.

#### 8.3.9 Transitivity

As visualized in Figure 21, we calculate the transitivity for each system and look at its distribution and correlation with the patient and the measured matched PBMC i.e., ELISpot values. We see that the distributions for both patients are quite similar. Patient one produces values that are skewed towards the lower quartile while patient two values are better distributed around the median. The combined data of both patients show a positive Pearson correlation coefficient, with no statistical significance, with the matched PBMCs. Patients one and two show a similar positive correlation. This confirms the observed correlation with the global clustering coefficient.

**Figure 21.**
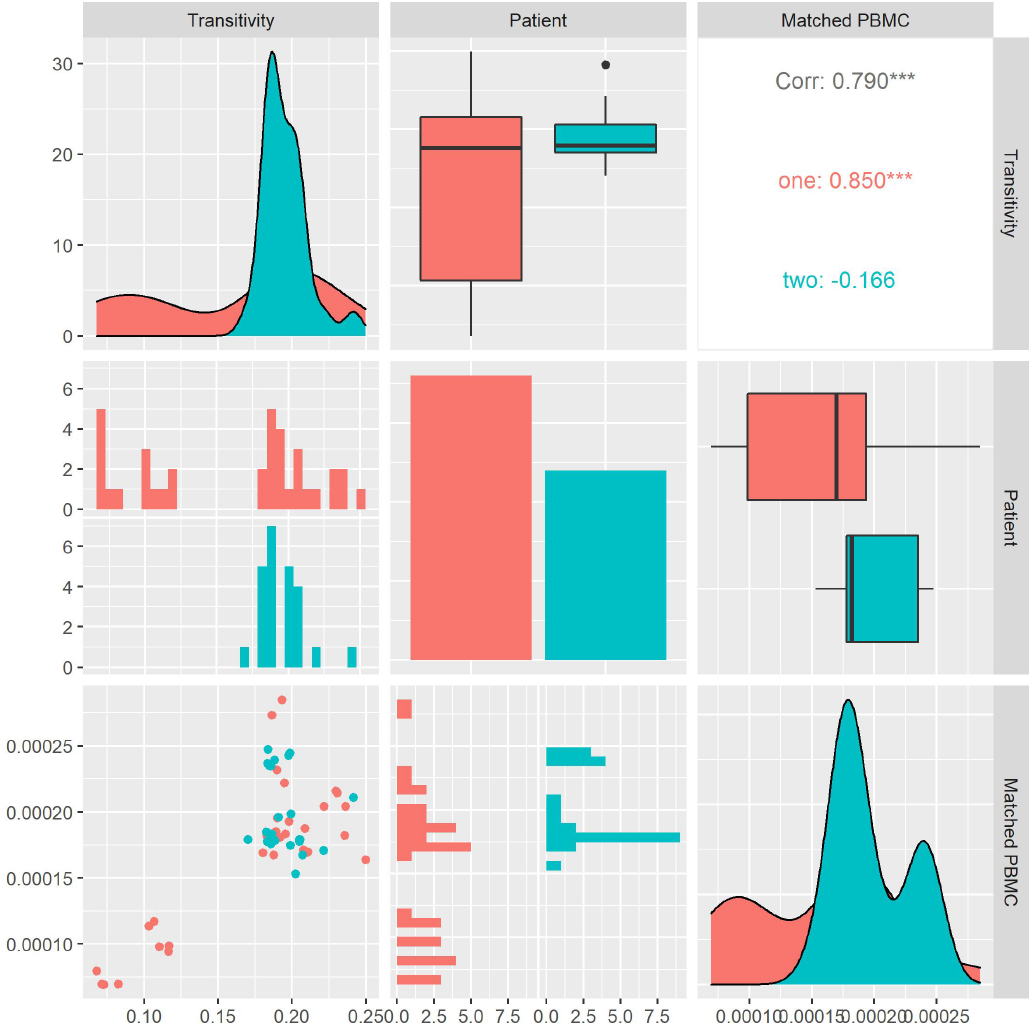
Patient Matched PBMCs vs. Transitivity

#### 8.3.10 Average Degree

As visualized in Figure 22, we calculate ⟨*k*⟩ for each system and look at its distribution and correlation with the patient and the measured matched PBMC i.e., ELISpot values. We see that the distributions for both patients are somewhat similar. Patient one produces values that are skewed towards the lower quartile while patient two values are better distributed around the median. The median of patient one is significantly higher. Combined data of both patients show an inverse Pearson correlation coefficient, with no statistical significance, with the matched PBMCs. Patient two shows a similar positive correlation in contrast to the patient one that shows a positive correlation. From these observations, we could imply that the impact of the density of the network on the matched PBMCs is patient-specific.

**Figure 22.**
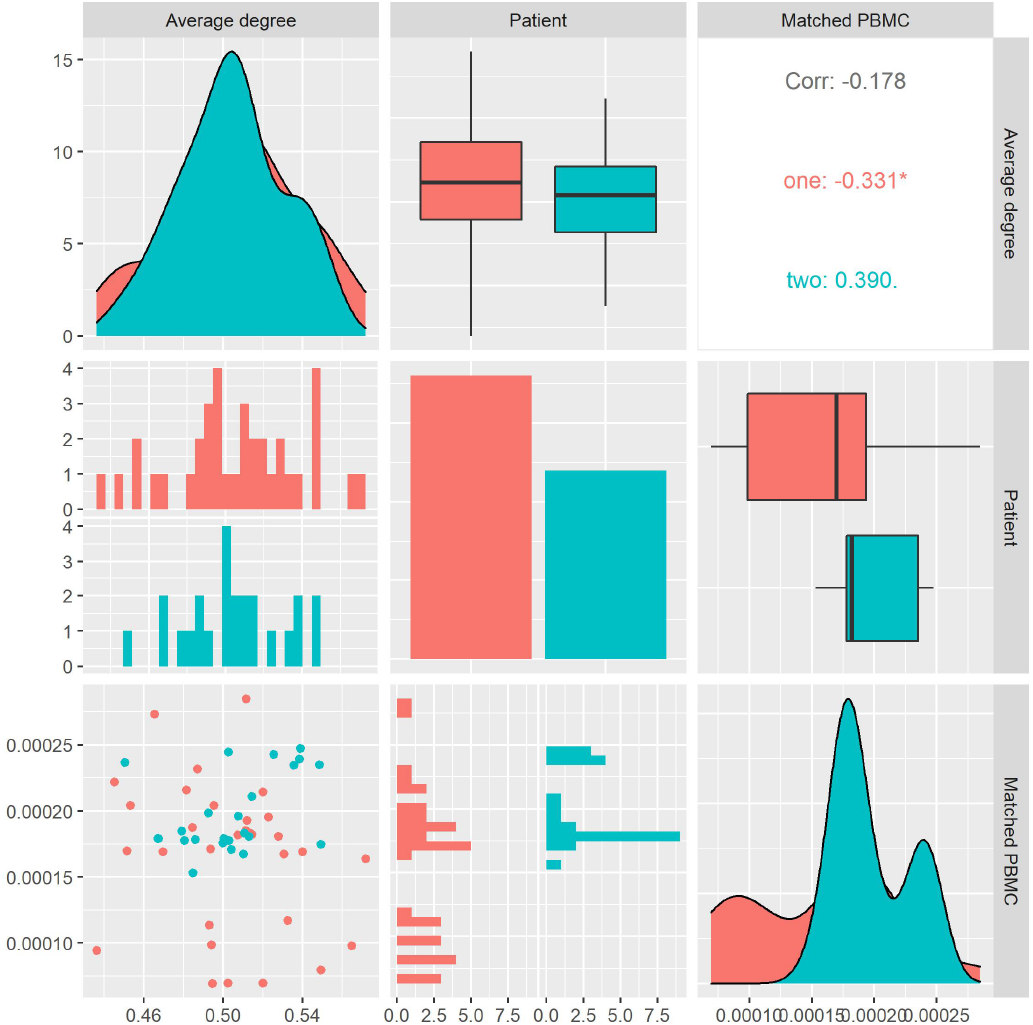
Patient Matched PBMCs vs. Average Degree

#### 8.3.11 Degree Centrality

As visualized in Figure 23, we calculate the degree centrality for each system and look at its distribution and correlation with the patient and the measured matched PBMC, i.e., ELISpot values. We see that the distributions for both patients are different. Patient one produces values that are mostly skewed toward the lower quartile while patient two is skewed toward the upper quartile. The median of patient one is significantly higher. Combined data of both patients show a positive Pearson correlation coefficient with a statistical significance indicated by a p-value < 0.05, with the matched PBMCs. Patient two shows a similar positive correlation in contrast to patient one, which shows a correlation near zero. From these observations, we could imply that the impact of a high number of incoming and outgoing edges from the vertices in the system has a positive impact on the matched PBMCs. There is, however, a patient-specific aspect in the magnitude of the positive impact.

**Figure 23.**
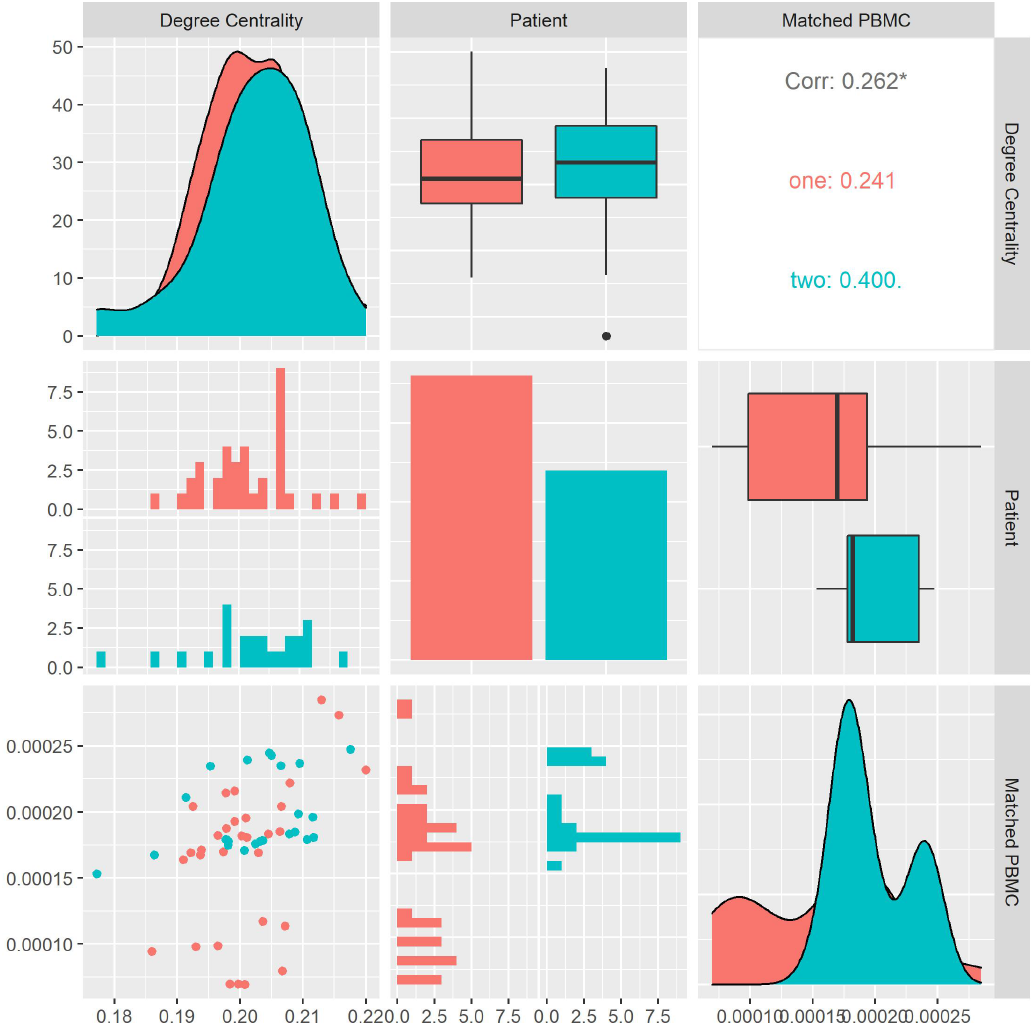
Patient Matched PBMC’s vs. Degree Centrality

#### 8.3.12 Average Core Hubs

As visualized in Figure 24, we calculate the average core hubs for each system and look at its distribution and correlation with the patient and the measured matched PBMC i.e., ELISpot values. We see that the distributions for both patients are different. Patient one produces values that are mostly skewed toward the lower quartile, while patient two is skewed toward the upper quartile. The median of patient one is significantly higher. Combined data of both patients show a positive Pearson correlation coefficient with statistical significance indicated by a p-value < 0.01, with the matched PBMCs. Patient two shows a similar positive correlation in contrast to patient one which shows a correlation near zero. From these observations, we could imply that the impact of a high number of incoming and outgoing edges from the vertices in the system has a positive impact on the matched PBMCs. There is however a patient-specific aspect in the magnitude of the positive impact. This observation agrees with the observations of Degree Centrality and its correlation with the matched PBMC values.

**Figure 24.**
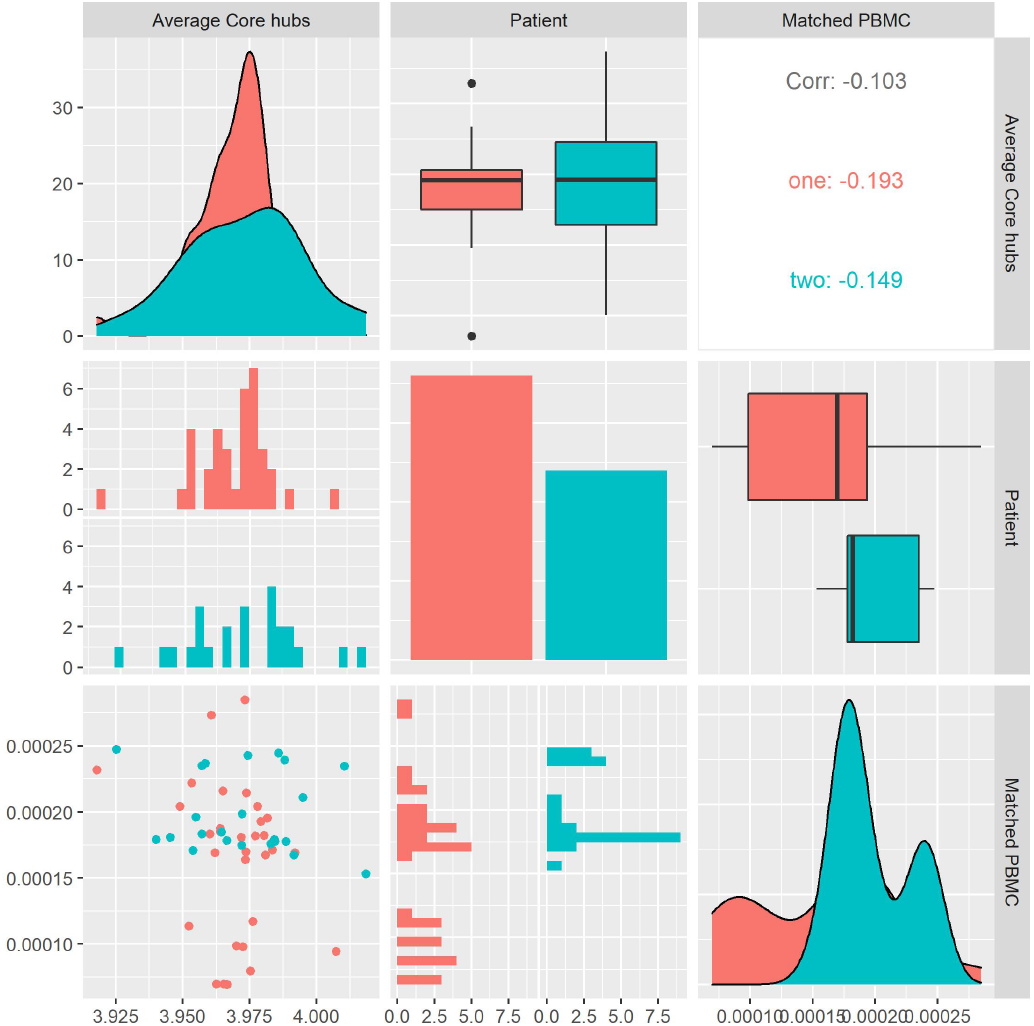
Patient Matched PBMCs vs. Average Core Hubs

#### 8.3.13 Density

As visualized in Figure 25, we calculate the density for each system and look at its distribution and correlation with the particular patient and the measured matched PBMCs i.e., ELISpot values. We see that the distributions for both patients are different. Patient one produces values that are mostly skewed toward the lower quartile while patient’s two is skewed toward the upper quartile. The median of patient one is significantly higher. Combined data of both patients show a positive Pearson correlation coefficient with statistical significance indicated by a p-value < 0.01, with the matched PBMCs. Patient two shows a similar positive correlation in contrast to patient one which shows a correlation near zero. From these observations, we could imply that a high number of incoming and outgoing edges from the system’s vertices positively impact the matched PBMCs. There is however a patient-specific aspect in the magnitude of the positive impact. This observation agrees with the observations of Degree Centrality and Average Core Hubs. As the measures, Density, Degree Centrality, and Average Core Hubs overlap it makes sense to select the one with the highest correlation and statistical significance i.e., Average Core Hubs.

**Figure 25.**
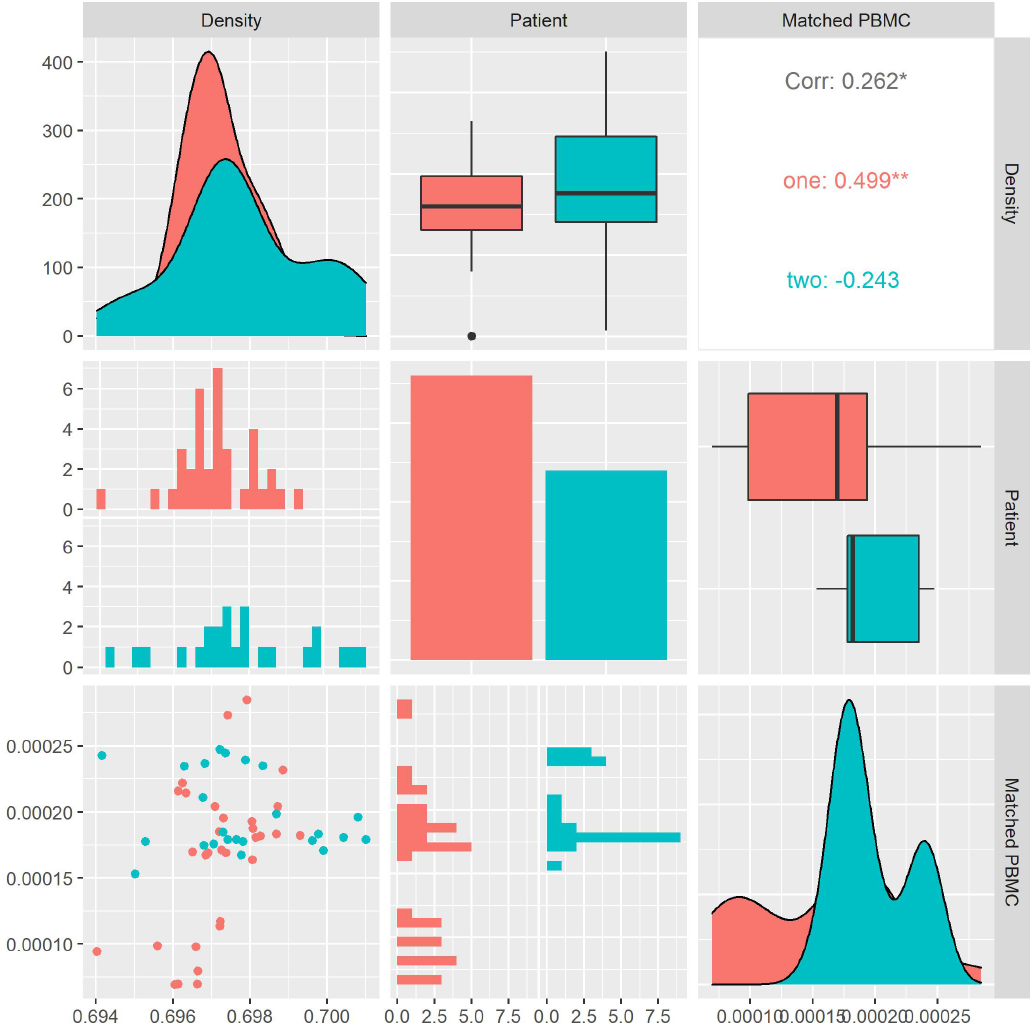
Patient Matched PBMC’s vs. Density

#### 8.3.14 Betweenness Centrality

As visualized in Figure 26, we calculate the Betweenness Centrality for each system and look at its distribution and correlation with the particular patient and the measured matched PBMC i.e., ELISpot values. We see that the distributions for both patients are different. Patient one produces values that are mostly skewed towards the lower quartile while patient’s two is skewed towards the upper quartile. The median of patient one is significantly higher. Combined data of both patients show a positive Pearson correlation coefficient with statistical significance indicated by a p-value < 0.01, with the matched PBMCs. Patient two shows a similar positive correlation in contrast to patient one which shows a correlation near zero. From these observations, we could imply that the impact of high number of nodes in a system that is on the path between other nodes has a positive impact on the matched PBMCs. There is however a patient-specific aspect in the magnitude of the positive impact.

**Figure 26.**
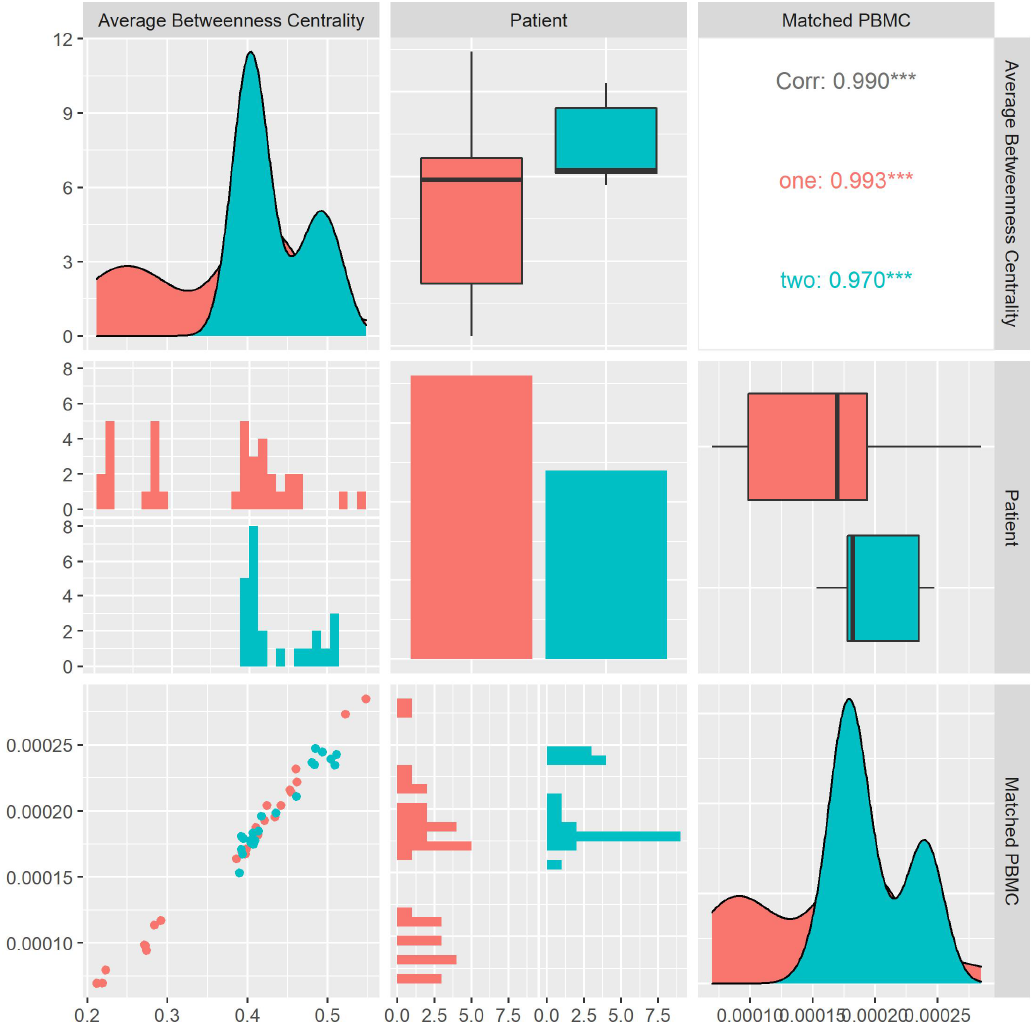
Patient Matched PBMC’s vs. Average Betweenness Centrality

#### 8.3.15 Closeness Centrality

As visualized in Figure 27, we calculate the Closeness Centrality for each system and look at its distribution and correlation with the particular patient and the measured matched PBMC i.e., ELISpot values. We see that the distributions for both patients are different. Patient one produces values that are mostly skewed towards the lower quartile while patient’s two are evenly distributed. The median of patient one is significantly higher. Combined data of both patients show a positive Pearson correlation coefficient with statistical significance indicated by a p-value < 0.05, with the matched PBMCs. Patient one and two shows a similar positive correlation. From these observations, we could imply that the impact of high number of nodes in a system with a high number of shortest distances to all other nodes, has a positive impact on the matched PBMCs.

**Figure 27.**
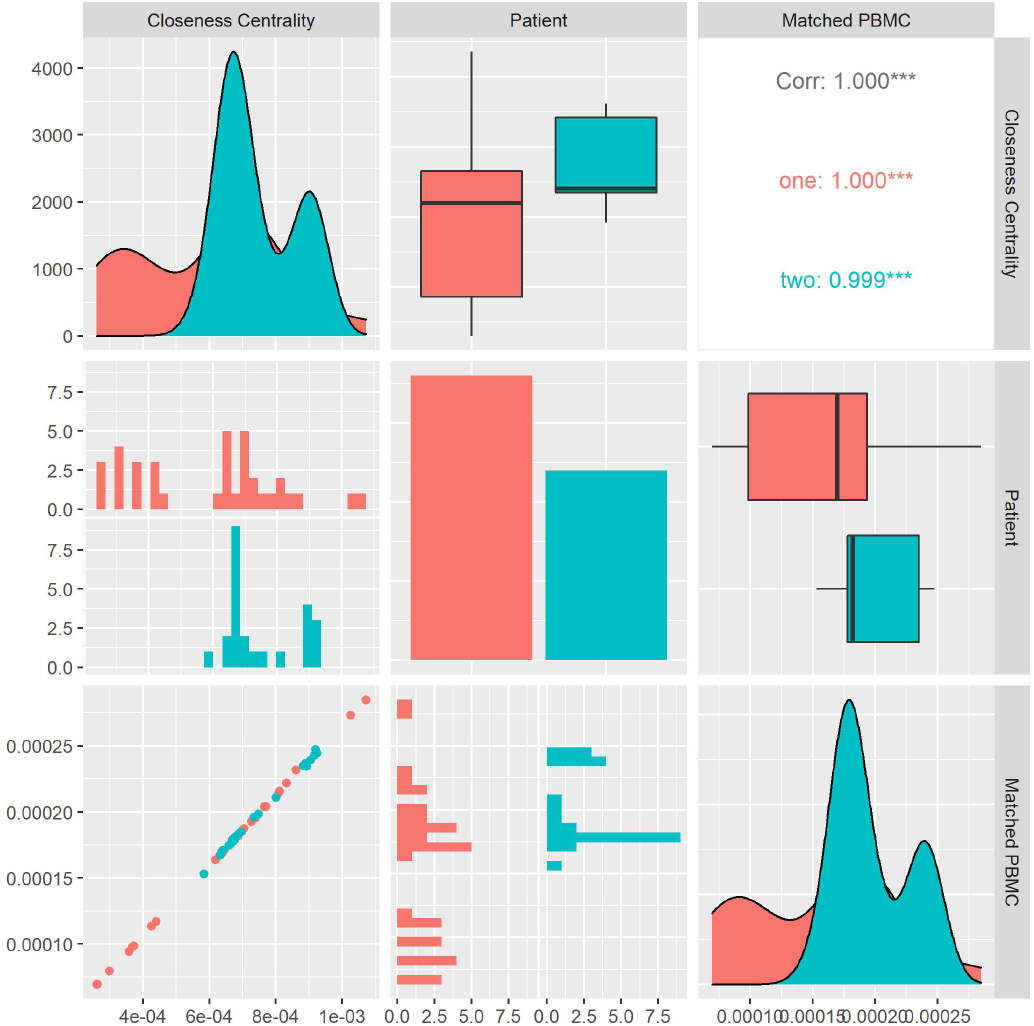
Patient Matched PBMC’s vs. Closeness Centrality

#### 8.3.16 Degree Pearson Correlation Coefficient

As visualized in Figure 28, we calculate the Degree Pearson correlation Coefficient for each system and look at its distribution and correlation with the particular patient and the measured matched PBMCs i.e., ELISpot values. We see that the distributions for both patients are different. Patient’s one and two produce values that are mostly skewed towards the upper quartile but there is a significant difference in their medians. Combined data of both patients show a positive Pearson correlation coefficient with statistical significance indicated by a p-value < 0.05, with the matched PBMCs. Patients one and two show a similar positive correlation. From these observations, we could imply that, a high number of similar connections between nodes in a system e.g., monomers, polymers, and atoms have a positive impact on the matched PBMCs.

**Figure 28.**
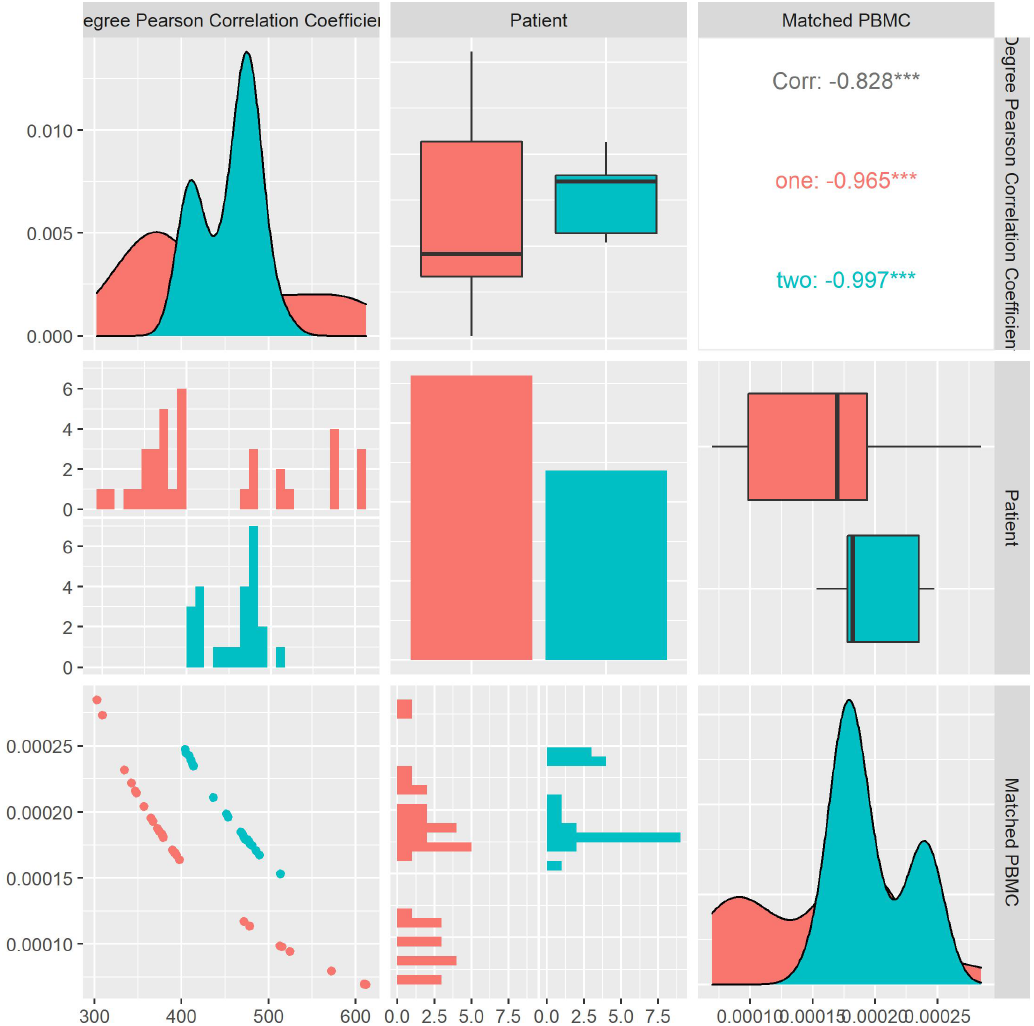
Patient Matched PBMC’s vs. Degree Pearson Correlation Coefficient

#### 8.3.17 Eigen Vector Centrality

As visualized in Figure 29, we calculate the average Eigen Vector Centrality for each system and look at its distribution and correlation with the particular patient and the measured matched PBMC i.e., ELISpot values. We see that the distributions for both patients are very similar. Patients one and two produce values that are mostly skewed towards the upper quartile with patient two having a longer tail. Combined data of both patients show a positive Pearson correlation coefficient with no statistical significance, with the matched PBMCs. However, patient two shows an inverse correlation in contrast to the positive correlation of patient one. From these observations, we could imply that the importance of this measure towards the impact on the matched PBMCs is not significant and it is patient-specific.

**Figure 29.**
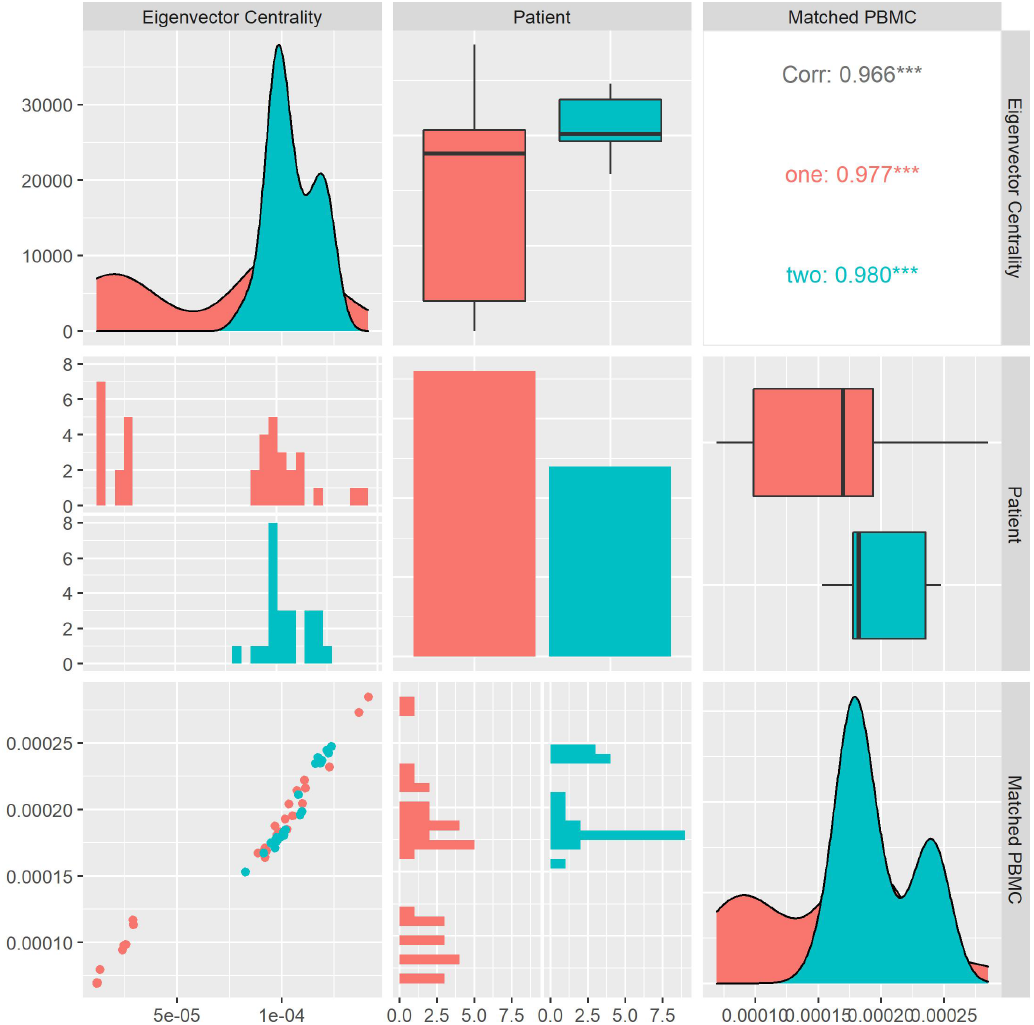
Patient Matched PBMC’s vs. Eigen Vector Centrality

#### 8.3.18 Page Rank Centrality

As visualized in Figure 30, we calculate the average Page Rank Centrality for each system and look at its distribution and correlation with the particular patient and the measured matched PBMC i.e., ELISpot values. We see that the distributions for both patients are very different. Patients one and two produce similar medians but patient one distribution is skewed toward the lower quartile and patient two’s toward the higher quartile. Combined data of both patients show a positive Pearson correlation coefficient with statistical significance shown by a p-value < 0.05, with the matched PBMCs. Patient two has a similar correlation. From these observations, we could imply that systems with a high number of nodes that have a high centrality based on connections from nodes that in their turn have high centrality have a positive influence on the produced PMBCs. However, this influence seems to be patient-specific.

**Figure 30.**
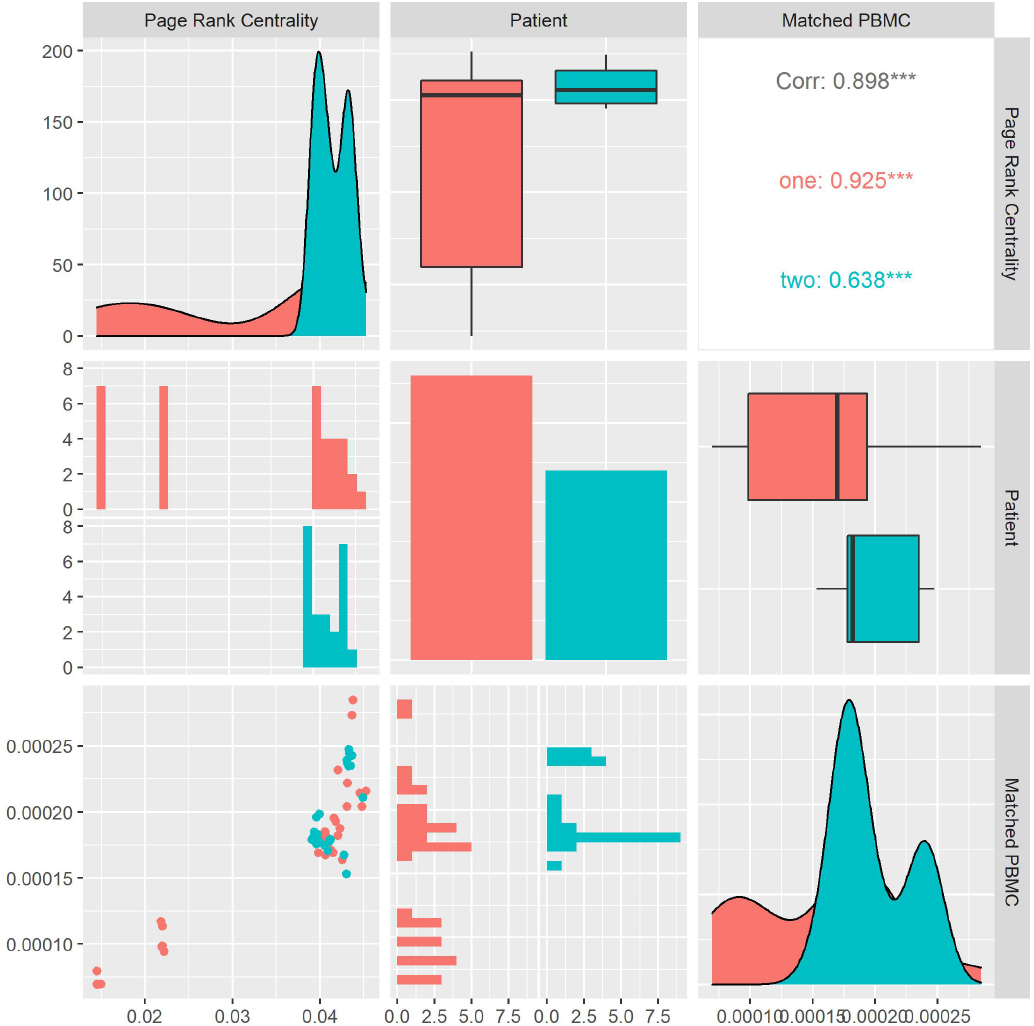
Patient Matched PBMC’s vs. Page Rank Centrality

#### 8.3.19 Attribute Assortativity Coefficient Source

As visualized in Figure 31, we calculate the average Attribute Assortativity Coefficient for the attribute Source for each system and look at its distribution and correlation with the particular patient and the measured matched PBMC i.e., ELISpot values. We see that the distributions for both patients are very different. Patients one and two produce similar medians but patient one distribution is skewed toward the upper quartile and patient two towards the lower quartile. Combined data of both patients show an inverse Pearson correlation coefficient with statistical significance shown by a p-value <0.05, with the matched PBMCs. Patients one and two have a similar correlation. From these observations, we could imply that systems with a more diverse unique node distribution have a higher likelihood to produce more PMBCs.

**Figure 31.**
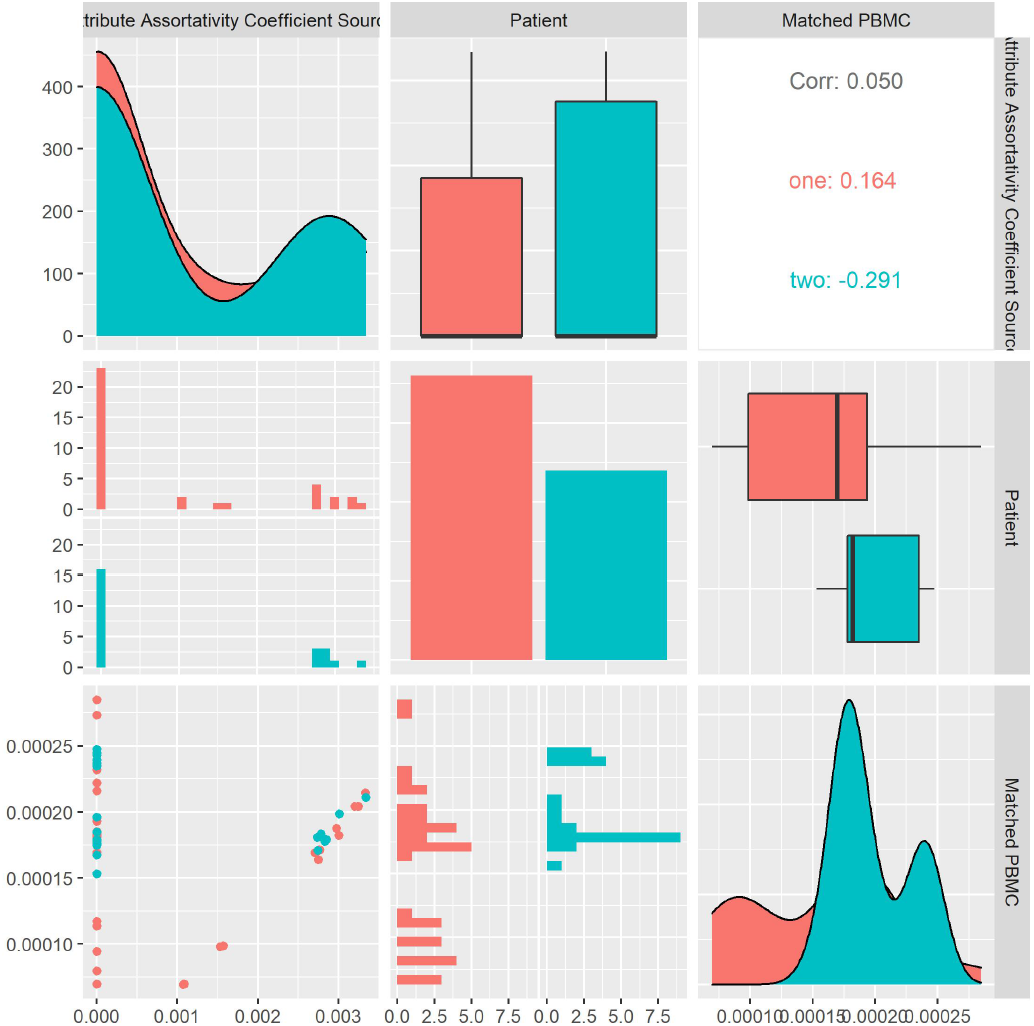
Patient Matched PBMC’s vs. Attribute Assortativity Coefficient Source

#### 8.3.20 Causality

A Bayesian network is a probabilistic model that represents the variables and their conditional dependencies as a directed acyclic graph (DAG). In a DAG, variables are represented as vertices and conditional dependencies as edges. Vertices that are not connected represent conditionally independent variables. Each node is associated with a probability function that takes in, a particular set of values for the node’s parent variables, and gives the probability, or probability distribution, of the variable represented by the node. As shown in Figure 32, we apply a structure learning algorithm to the data to learn the structure of the directed acyclic graph (DAG) from the data. Concretely we apply Incremental Association (IAMB) algorithm, based on the Markov blanket detection algorithm of the same name, which is based on a two-phase selection scheme (a forward selection followed by an attempt to remove false positives). Analyzing the DAG, we see that the matched PBMC, vectors, Average core hubs, Attribute Assortativity Coefficient Source, Size largest component, Eigen vector Centrality, and closeness centrality are conditionally dependent on the patient. This observation confirms observations made the correlation analysis. This indicates that system structures have patient-specific attributes and their matched PBMCs are as well patient specific.

**Figure 32.**
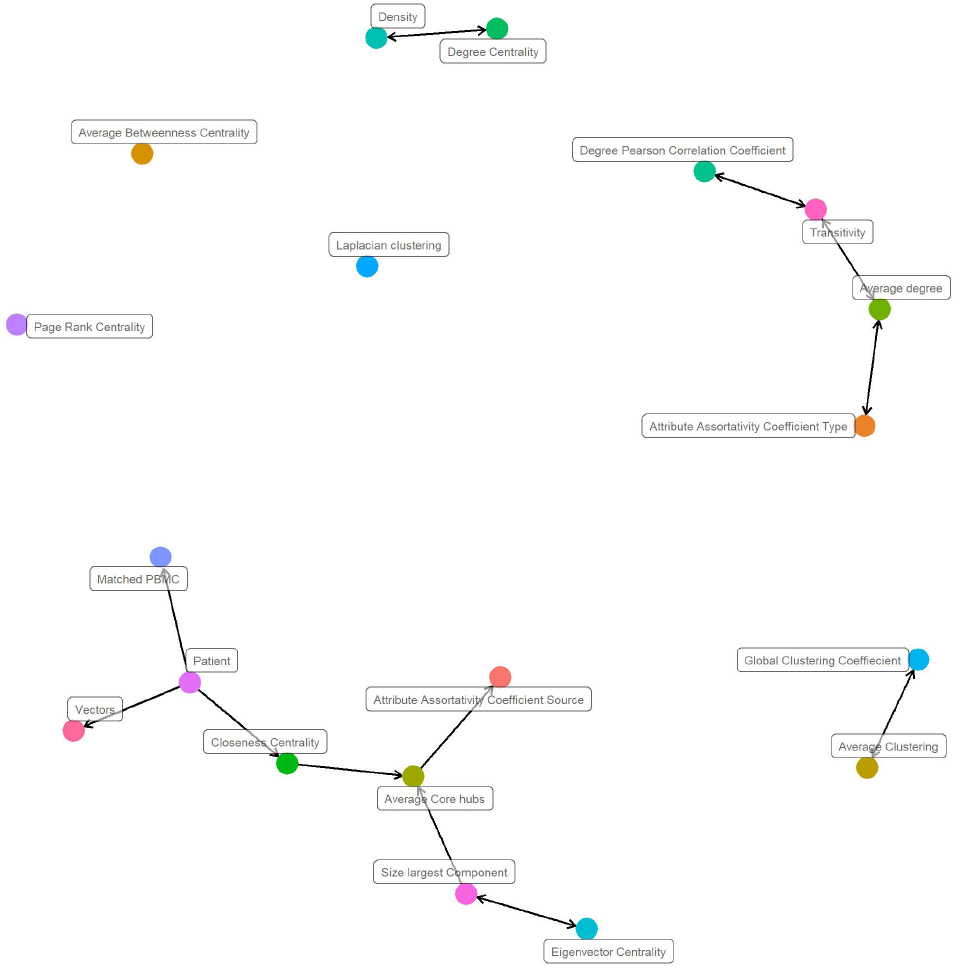
Structure learning casual network.

#### 8.3.21 Personal Models cross evaluation

We evaluate the performance of the personal model, trained on data from patient one, on data from patient two, and vice versa. As shown in Table 2, the metrics Precision, Recall, and F-measure drop significantly in contrast to their respective values in Table 2 and Table 3. We can conclude that the models do not necessarily generalize to unseen data from other patients and that personalized models perform better.

### 8.4 Evaluation metrics

#### 8.4.1 Accuracy

Accuracy is the most intuitive performance measure and it is simply a ratio of correctly predicted observations to the total observations.

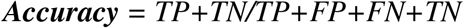

#### 8.4.2 Precision

Precision is the ratio of correctly predicted positive observations to the total predicted positive observations.

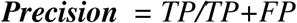

#### 8.4.3 Recall

Recall (Sensitivity) is the ratio of correctly predicted positive observations to all observations in actual class – yes

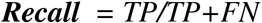

#### 8.4.4 F-measure

F-measure is the weighted average of Precision and Recall. Therefore, this score takes both false positives and false negatives into account. Intuitively it is not as easy to understand as accuracy, but F1 is usually more useful than accuracy, especially if you have an uneven class distribution. Accuracy works best if false positives and false negatives have similar cost. If the cost of false positives and false negatives are quite different, it’s better to look at both Precision and Recall.

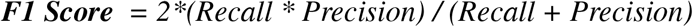

## AUTHOR CONTRIBUTIONS

IJ performed the statistical analysis and wrote the first draft of the manuscript. IJ and NM contributed to the conception and design of the study. LM and JM contributed to the medical aspect of the study. All authors contributed to the manuscript revision and read and approved the submitted version.

## DATA AVAILABILITY STATEMENT

The datasets for this study can be found in the In silico Antibody-Peptide Epitope prediction for Personalized cancer therapy.

## TRAINED MODELS AVAILABILITY STATEMENT

The trained models from this study will be made available for usage on Linked Chain website under the section A.I. tools.

## PERMISSION TO REUSE AND COPYRIGHT

Figures, tables, and images will be published under a Creative Commons CC-BY licence and permission must be obtained for use of copyrighted material from other sources (including republished/adapted/modified/partial figures and images from the internet). It is the responsibility of the authors to acquire the licenses, to follow any citation instructions requested by third-party rights holders, and cover any supplementary charges.

